# Photothermal transport for guiding nanoparticles through the vitreous humor

**DOI:** 10.1101/2025.07.24.666559

**Authors:** Léa Guerassimoff, Yera Ussembayev, Louise De Clerck, Deep Punj, Martijn van den Broek, Filip Beunis, Katrien Remaut, Kevin Braeckmans, Stefaan C. De Smedt, Félix Sauvage

## Abstract

Visual impairments affect over 2.2 billion people worldwide, yet delivering drugs to the eye’s posterior segment, including the retina, remains a major challenge. Intravitreal injection, the standard administration route to the posterior segment of the eye, often results in suboptimal drug diffusion through the vitreous, preventing drugs from reaching the retina. While various strategies have been explored to enhance the mobility of drug molecules and nanomedicines (drugs encapsulated in nanoparticles) in the vitreous, no method has demonstrated ‘guided transport’ of drugs and particles through the vitreous. In this study, we explore photothermal transport of nanoparticles in the vitreous using a pulsed-laser and indocyanine green added to the vitreous, both being clinically approved modalities. We found that photothermal transport allows to guide nanoparticles from one place in the vitreous towards the laser illuminated area in the vitreous, at a distance of the injection spot of the nanoparticles. Using multiple-particle tracking and numerical simulations, we found that both thermal convection and thermophoresis contribute to photothermal transport of nanoparticles in the vitreous. We identified parameters for optimizing this effect, including dye concentration, particle size, distance from the laser focus, and laser fluence. Our findings establish a novel and clinically relevant paradigm for light-guided drug delivery in the eye. This study represents, to our knowledge, the first demonstration of guided light-controlled particle transport in the vitreous using ocular dyes and pulsed-lasers which are routinely used in ophthalmology.

## Introduction

Visual impairments affect more than 2.2 billion of people worldwide, with refractive errors (RE), cataract (CAT), diabetic retinopathies (DR), glaucoma (GL), and age-related macular degeneration (AMD) as most frequent disorders.^1,2^ For many eye diseases, successful treatment remains a significant challenge. This is partially explained by the fact that drug delivery to specific compartments in the eye is still extremely difficult^3^ due to the presence of various ocular barriers (e.g. lens, corneal barriers, vitreous humor, inner limiting membrane (ILM)) and poor control over the location where the drug ends up following administration. The main drug administration route to treat the posterior segment of the eye (including the retina) is intravitreal (IVT) injection, being the injection of drugs into the vitreous humor. As an example, anti-VEGF antibodies are routinely intravitreally-injected for the treatment of AMD.^3,4^ However, there is still an unmet need for delivery strategies which allow ‘modern’ medication, including antibodies and nanomedicines, to precisely reach the retina. Critical challenges persist following IVT injections, particularly in ensuring adequate drug diffusion through the vitreous humor and effective accumulation in the retina.

The vitreous humor is a gel-like structure^5,6^, filling the space between the lens and the retina, which is considered a key structural component of the eye as it helps maintain its shape and protects it from mechanical damage.^7^ When the eye has reached its adult size (typically between 14 and 18 years), this transparent and highly-hydrated extracellular matrix is composed of two phases:^8^ (i) a bulk gel part representing 80 % of the total volume of the vitreous humor, and (ii) a liquid phase.^9^ The vitreous mainly consists of water (98 to 99 vol %)^7,10^ and a network of biomacromolecules composed of collagen fibers, hyaluronan and proteoglycans.^5^ This 3D collagen network, which has been characterized by a mesh diameter of around 500 nm^9,11^, significantly contributes to the viscosity of the vitreous humor. Indeed, collagen fibers can vary in density, length and texture, as can the ratio of collagen to hyaluronan, both of which significantly affect the viscosity of the vitreous.^9^

As a dynamic barrier, the vitreous humor undergoes partial liquefaction with aging, changing its physicochemical and mechanical properties.^12^ Four different states have been identified^12^: (i) an initial phase in which the vitreous remains a homogeneous gel-like substance; (ii) the onset of liquefaction characterized by progressive dehydration of the gel and formation of isolated liquid pockets; (iii) extensive liquefaction during which these pockets enlarge and coalesce; and (iv) post vitreous detachment (PVD) which occurs as a result of vitreous liquefaction combined with the weakening of the vitreoretinal adhesion. While barely studied as of today, structural changes in the aging vitreous may significantly influence drug distribution following IVT injection, as the drug molecules may tend to follow connective flow patterns within the liquefied regions.^9^ Moreover, the dense macromolecular matrix of the vitreous hinders diffusion, particularly for larger molecules (such as biotherapeutics) and nanomedicines.^11,13^ Additionally, drug reflux^1^ and ‘multi-directional diffusion’ away from the injection spot in the vitreous site may further compromise drug delivery, substantially reducing the likelihood of the drug reaching the retina.

Developing innovative strategies to enhance drug accumulation and penetration into the retina following IVT injection remains a significant challenge while such strategies could significantly help in retinal disease management. Improvements in retinal drug delivery might be achieved through two main strategies. One strategy is the design of (drug loaded) nanocarriers with surface properties that avoid their aggregation in the vitreous (which prevents them from reaching the retina); examples include micelles^14^, liposomes^15^ and lipid nanoparticles^16^. Another strategy is based on the application of external stimuli into the vitreous to actively ‘guide’ drugs from the injection spot in the vitreous towards the retina. Delivering IVT-injected therapeutics to a specific region of the retina based on external stimuli has, however, rarely been reported. One study reports on light-driven nitrogen-doped TiO_2_ nanomotors^17^ moving in vitreous while another study describes helical magnetic micro-propellers whose motion in vitreous can be controlled by a rotating magnetic field.^18^ While these are praiseworthy efforts, the design and synthesis of micro-propellers and nanomotors remains rather complex, while their ocular safety remains unclear. Therefore, exploring strategies that could safely enable motion of molecules (like biotherapeutics) and nanoparticles (like nanomedicines) in the vitreous, using materials and external stimuli approved for clinical use in the vitreous remains highly attractive.

Photothermal transport is the phenomenon in which molecules or particles in a medium (fluid) move due to a local temperature increase of the medium induced by light absorption. This motion results from a combination of mechanisms, the primary ones being (i) thermal convection, (ii) Marangoni effect (i.e., mass transfer along an interface between two phases due to a gradient of the surface tension) and (iii) thermophoresis.^19–21^ Thermal convection^22^ occurs when heat causes density variations in a fluid. When the temperature of the fluid near a laser-induced hot spot increases, the density lowers locally. This warmer, less dense, fluid becomes buoyant and rises upward while the surrounding cooler (and denser) fluid moves downward. This results in a circulating continuous fluid motion, known as thermal convection, around the heated region. Unlike photothermal convection, opto-thermophoresis^23^ does not involve a bulk fluid motion but a *directed* force acting on individual molecules/particles: the temperature gradient *around* each molecules/particle creates an imbalance in local intermolecular interactions which drives the molecules/particles either toward or away from the heat source. This force has been explored to manipulate various objects like cells^24^, particles^25^ and DNA^26^ in media like water^27^ and cyclohexane^28^. Photothermal convection and opto-thermophoresis have been mostly studied to enable motion of molecules/particles on the surface of a light absorbing (solid) substrate (e.g. a plasmonic or metal surface)^27,29^ which is, however, challenging to realize *in vivo*. In the current study we aim to explore the potential of light absorbing dyes in solution as heating substrates for the induction of photothermal transport of nanoparticles (NPs) suspended in the solution. Next we aim to elucidate if photothermal transport of nanoparticles can be achieved in the vitreous, using dyes and lasers which are in current ophthalmological use. More specifically we explore the potential of nanosecond pulsed-lasers and vital dyes, such as indocyanine green (ICG) and trypan blue (TB), that are commonly used in the clinic for the staining of ocular tissues^30,31^ and for retinal angiography.^32,33^ We aim to provide a sound physical understanding of particle motion driven by photothermal transport in the vitreous, investigate this phenomenon experimentally using multiple-particle tracking (MPT), and aim to reveal whether directed (‘guided’) transport of nanoparticles in the vitreous based on opto-thermophoresis and thermal convection can be achieved (**Figure 1**). To the best of our knowledge, photothermal transport in biological matrices has never been reported. Also, while vital dyes and pulsed-lasers are widely used in ocular interventions, their combined use in the eye is uncommon. The current study is part of our efforts^34^ to widely explore the potential of a combined use of ocular dyes and pulsed-lasers for novel treatments of eye diseases and advanced ocular surgeries.

**Figure 1.**
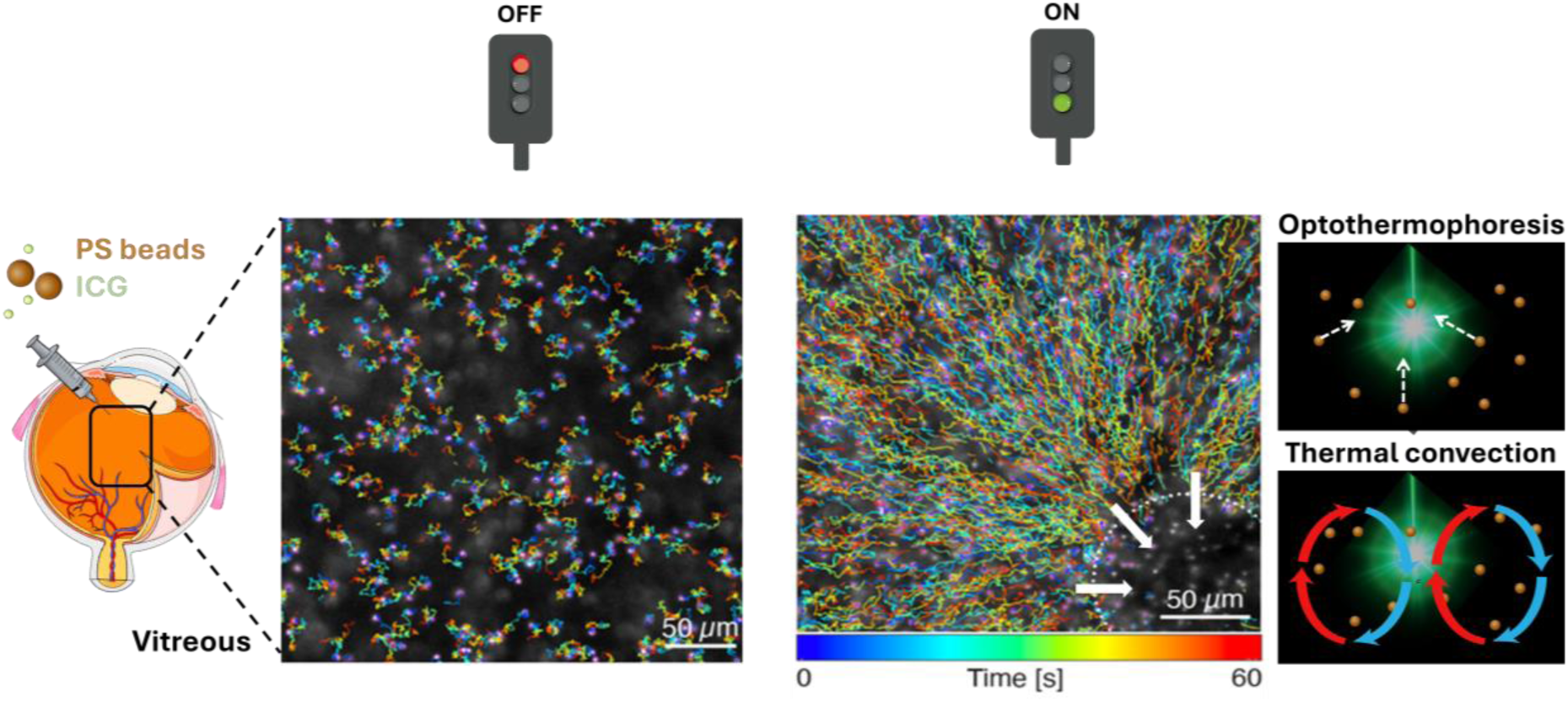
Schematic representation of photothermal transports, opto-thermophoresis and thermal convection, to guide polystyrene (PS) nanoparticles through vitreous. PS nanoparticles and the ocular dye indocyanine green (ICG) are added to the medium (water or vitreous) which is locally irradiated with a nanosecond pulsed-laser. The left panel shows the random Brownian motion of nanoparticles when the laser is off. The middle panel shows the directed motion of nanoparticles when the laser is on: nanoparticles moved towards the laser illuminated spot (indicated by the white circle). The right panel shows the two main mechanisms involved in the photothermal transport of nanoparticles through the vitreous: optothermophoresis (top panel), the white arrows illustrate the direction of the motion of nanoparticles, and thermal convection (bottom panel).

## Results

### Photothermal transport of polystyrene (PS) nanoparticles in water

Fluorescent PS nanoparticles (520 nm) were suspended (dilution ratio of 1/1000 (v/v)) in aqueous solutions of indocyanine green (ICG, 0.5 mg/mL), as illustrated in **Figure 2A**. After depositing droplets of these mixtures on microscope slides, selected areas within the droplets were irradiated with nanosecond laser pulses (532 nm; pulse duration 2-5 ns, 100 Hz repetition rate at a pulse fluence of 2.07 J.cm^-2^) (**Figure 2B**) and videos were recorded (1 fps, before (15 s), during (30 s) and after (15 s) laser irradiation) (see **Supplementary Table 1**). This initial experiment revealed the attraction of nanoparticles toward the laser beam focus **(Video S1**) and their subsequent accumulation within the illuminated zone (**see Supplementary Figure S1A**). Interestingly, no attraction / accumulation of PS nanoparticles towards / at the laser focus was observed in the absence of ICG (**Video S2**) or without laser irradiation (**Video S3**). This shows that photothermal particle motion is dependent on both the presence of the dye and laser exposure. Note that, in both control conditions (i.e., without ICG or without laser irradiation), PS nanoparticles still exhibited Brownian motion (**Figure 2C**).

**Figure 2.**
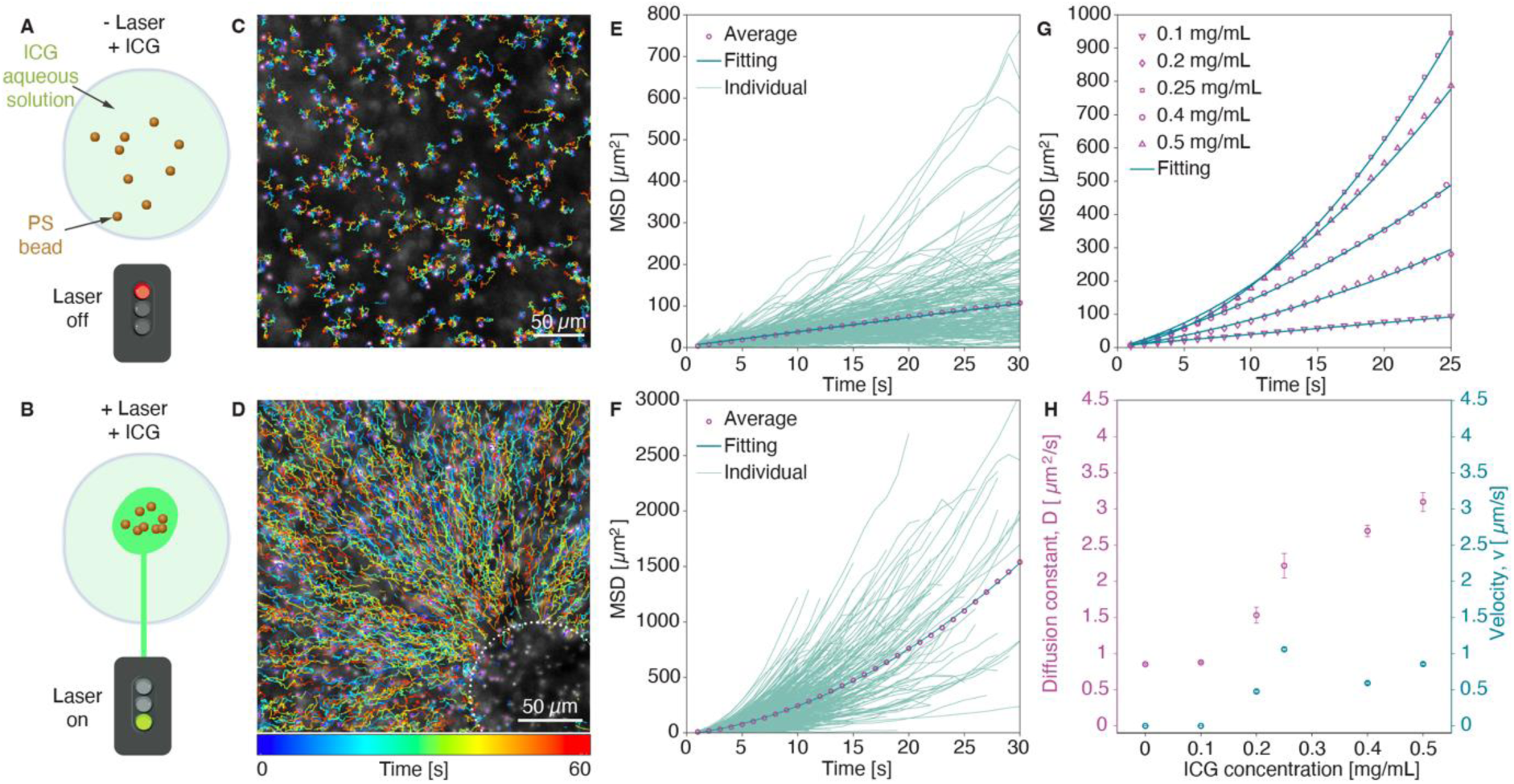
Photothermal transport of PS nanoparticles (dp = 520 nm) in the presence of ICG upon laser irradiation in water. **A**: Schematic illustration of PS nanoparticles mixed with ICG in water without laser exposure. **B**: Schematic representation of PS nanoparticle transport under laser irradiation. **C**: Representation of colored single traces of PS nanoparticles (dilution ratio 1/1000 (v/v)) from MPT analysis in water samples in the presence of ICG at 0.5 mg/mL and without laser irradiation (**Video S3**). **D**: Representation of colored single traces of PS nanoparticles (dilution ratio 1/1000 v/v) from MPT analysis in water samples in the presence of ICG at 0.5 mg/mL and upon laser irradiation (2.07 J/cm^2^, 532 nm). The laser spot is located within the white circle. (**Video S1**). **E**: Evolution of MSD values as a function of time obtained from MPT analysis of **Video S3** (i.e., in water, PS nanoparticles at a dilution ratio of 1/1000 (v/v), ICG at 0.5 mg/mL, no laser irradiation). Representation of individual nanoparticles traces and implementation of a linear fitting model (diffusion regime). Number of traces for analysis = 609. **F**: Evolution of MSD values as a function of time obtained from MPT analysis of **Video S1** (i.e., in water, PS nanoparticles at a dilution ratio of 1/1000 (v/v), ICG at 0.5 mg/mL, upon laser irradiation (2.07 J/cm^2^, 532 nm)). Representation of individual particles traces and implementation of a quadratic fitting model. Number of traces for analysis = 2326. **G**: Time traces of average MSD acquired for different ICG concentrations (i.e., 0, 0.1, 0.2, 0.25, 0.4 and 0.5 mg/mL) in water (PS nanoparticles at a dilution ratio of 1/5000 (v/v)) from the extracted individual time traces summarized in **Supplementary Figure S2**. **H**: Evolution of extracted fitting parameters, the diffusion constant *D* (left Y axis) and the particle velocity *v* (right Y axis), as a function of ICG concentration (i.e., 0, 0.1, 0.2, 0.25, 0.4 and 0.5 mg/mL) in water (PS nanoparticles at a dilution ratio of 1/5000 (v/v)).

To characterize the nature of particle motion, we recorded time lapse videos and applied multi-particle tracking (MPT) analysis^11,35–37^ (**Videos S1-3**). According to Einstein’s relation, the diffusion coefficient *D* is given by:

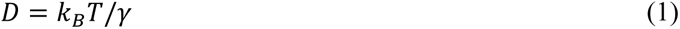

where *k_B_* = 1.38×10^-23^ J/K is the Boltzmann constant, *T* is the absolute temperature, and *γ* is the friction coefficient experienced by the particle in the fluid medium. For spherical particles, the friction coefficient *γ*, also known as the hydrodynamic drag, is related to the particle diameter *d_p_* and the temperature-dependent dynamic viscosity *η* of the liquid by Stokes’ law:

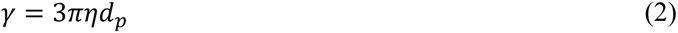

To quantify particle trajectories, we tracked (see **Methods**) the two-dimensional (2D) positions (*x*, *y*) of individual particles across time *t* from video frames (**Figures 2C-D**). From this tracking data, the mean-square displacement (MSD) as a function of lag time *τ* was calculated as:

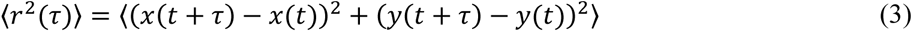

The MSD provides insights into the nature of particle motion - whether purely diffusive or influenced by an external force. In control experiments (i.e., **Video S2**: no ICG, no laser; and **Video S3**: ICG, no laser), individual particle tracks display random, non-directional Brownian motion (**Figure 2C**). This is confirmed by the MSD profile (**Figure 2E**) from **Video S3**, which fits well to a linear model for normal 2D diffusion^35^:

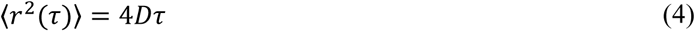

From this fit, the experimental diffusion constant was calculated as *D*_exp_ = 0.853 ± 0.007 µm^2^/s, closely matching the theoretical value *D*_th_ = 0.825 µm^2^/s for particles of 520 nm diffusing in water at room temperature. In contrast, under laser irradiation and in the presence of ICG (**Video S1, Supplementary Table 1**), particle trajectories (**Figure 2D**) exhibit directed motion toward the laser focal point. The corresponding MSD (**Figure 2F**) deviates from the linear model and instead follows a quadratic dependence, indicative of active transport^35^:

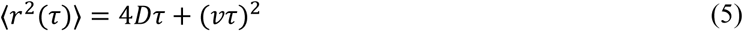

Here, *v* represents the velocity corresponding to the directed motion of the particles. Fitting this model to MSD data yields a significantly higher apparent diffusion constant *D*_exp_ = 2.68 ± 0.12 µm^2^/s and a velocity *v* = 1.160 ± 0.021 µm/s, demonstrating the influence of the photo-induced driving force on the motion of nanoparticles. Notably, the diffusion constant under laser irradiation is approximately 3 times higher than in control experiments (**Video S3**: ICG, no laser), due to the temperature rise and corresponding reduction in viscosity caused by the ICG absorption.

To gain deeper insight into the role of ICG in mediating photothermal transport of nanoparticles, we investigated a range of ICG concentrations: 0.1, 0.2, 0.25, 0.4, and 0.5 mg/mL, while keeping the PS particle dilution ratio constant at 1:5000 (v/v). The extracted individual time traces are shown in **Supplementary Figure S2** whereas the averaged MSD values are summarized in **Figure 2G**. Under pulsed laser irradiation (2.07 J/cm², 532 nm), PS particles exhibit enhanced motion toward the laser focal point at high ICG concentrations (i.e., 0.25-0.5 mg/mL), in contrast to the slower or negligible motion at lower concentrations (i.e., 0.1-0.2 mg/mL; see **Video S4** for 0.1 mg/mL). This trend is quantitatively illustrated in **Figure 2H** where the extracted parameters from MSD analysis, including the diffusion constant *D* and particle velocity *v*, are shown as a function of ICG concentration. At concentrations below 0.2 mg/mL, the particle motion remains purely Brownian motion. However, starting from 0.2 mg/mL, we observe additional directed motion. Above this threshold, both diffusion and particle velocity increase, with particle velocities reaching up to about 1 µm/s. Based on the measured diffusion constants and equations (1)-(2), we estimated the corresponding localized temperature (see **Methods**) increases for ICG concentrations of 0.1 and 0.5 mg/mL, which yields approximate temperatures of 20 °C and 85 °C, respectively. These results highlight the critical role of ICG in enabling laser-induced photothermal transport of PS nanoparticles through water, along with the ability to tune particle velocity by simply varying the dye concentration.

### Photothermal transport of PS particles in bovine vitreous

As a biphasic solid-like material, the vitreous humor consists of 80 % bulk gel while the remaining 20 % is liquid.^9^ The vitreous also contains a complex network of biomacromolecules including various types of collagen (e.g., II, V, XI and VI) and hyaluronan which all contribute to gel swelling by drawing liquid into this network.^7^ It is of great interest to investigate and characterize the motion of PS nanoparticles (i.e., MSD regime, diffusion constant and velocity) upon laser irradiation in the vitreous, as it could provide essential insights into the potential of light to move and guide intravitreally-injected nanomedicines. As reported below, the influence of the laser fluence on photothermal transport of PS nanoparticles in bovine vitreous was investigated along with the impact of the PS particle diameter and the distance of the PS particles from the laser focus. Moreover, thermal effects on the vitreous itself were investigated.

Fluorescent 520 nm PS nanoparticles were mixed (dilution ratio of 1/1000 (v/v)) with aqueous ICG solutions (0.5 mg/mL). The mixture of ICG and PS nanoparticles (50 µL) was then injected into 500 µL-vitreous samples, freshly excised and extracted from cow eyes, as illustrated in **Supplementary Figure S3**. Selected areas within the excised vitreous samples were first irradiated with a nanosecond pulsed-laser (532 nm) with a fluence ranging from 0.34 to 1.03 J/cm^2^ and videos were recorded (1 fps, 30 to 60 s) before (15 s), during (30 s) and after (15 s) laser irradiation (see **Supplementary Table 1**). This experiment in the vitreous first confirmed the attraction of the nanoparticles toward the laser beam and their subsequent accumulation in the illuminated zone (**Supplementary Figure S1B**, **Video S5, 0.69 J/cm²**). Control samples (i.e., without laser irradiation (**Video S6, 0.69 J/cm²**) or without ICG (**Video S7, 0.69 J/cm²**)) confirmed that the motion of particles was dependent on both the dye and light, as no particle attraction was observed in samples lacking ICG or laser irradiation.

Figure 3 presents multi-particle tracking analysis of PS nanoparticles in bovine vitreous, highlighting two distinct regimes of particle motion depending on laser irradiation conditions, as demonstrated in **Videos S5** and **S6**.

(i) **confined motion without laser irradiation (0 J/cm²):** in the absence of laser exposure (**Video S6**), PS nanoparticles exhibit confined, low-mobility behavior. Individual particle trajectories appear confined in space and randomly oriented (Figure 3A), consistent with restricted Brownian motion. The mean square displacement curve (Figure 3E) shows a saturating trend, indicative of particle confinement. This behavior is described using an exponential confinement equation^35^:

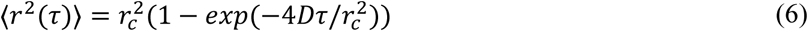

where *r_c_* is a confinement radius, defined as the characteristic spatial limit within which the particle remains trapped due to physical or environmental constraints (e.g., the mesh structure of the vitreous). It represents the maximum spatial extent of free diffusion before encountering restrictions. Fitting the expression (6) to the experimental data from **Video S6** yields *r_c_* = 0.77 ± 0.06 µm and a diffusion constant *D*_exp_ = 0.029 ± 0.006 µm^2^/s, significantly lower than the expected diffusion coefficient in water (0.825 µm^2^/s).
(ii) **directed motion with laser irradiation (0.34–1.03 J/cm²):** upon laser illumination (**Video S5**), PS nanoparticles exhibit clear directed motion toward the laser focus, forming elongated trajectories (Figures 3B**–D**) that become increasingly elongated with increasing laser fluence. The average MSD plots in Figure 3F are extracted from the individual traces shown in **Supplementary Figure S4** and display a pronounced quadratic time dependence, indicative of active transport. Figure 3F shows MSD curves for three laser fluence levels (0.34, 0.69, and 1.03 J/cm²), with steeper curves at higher fluences, reflecting faster directed motion. Figure 3G quantifies this trend, plotting the diffusion constants and velocities as a function of laser fluence. While the diffusion coefficient remains nearly constant, the particle velocity increases monotonically with the fluence, confirming the fluence-dependent nature of laser-induced transport (**Table 1**). Therefore, we have successfully demonstrated the photothermal transport of 520 nm-nanoparticles in the presence of ICG in the vitreous humor, as well as its strong dependency on the laser fluence value.

**Figure 3.**
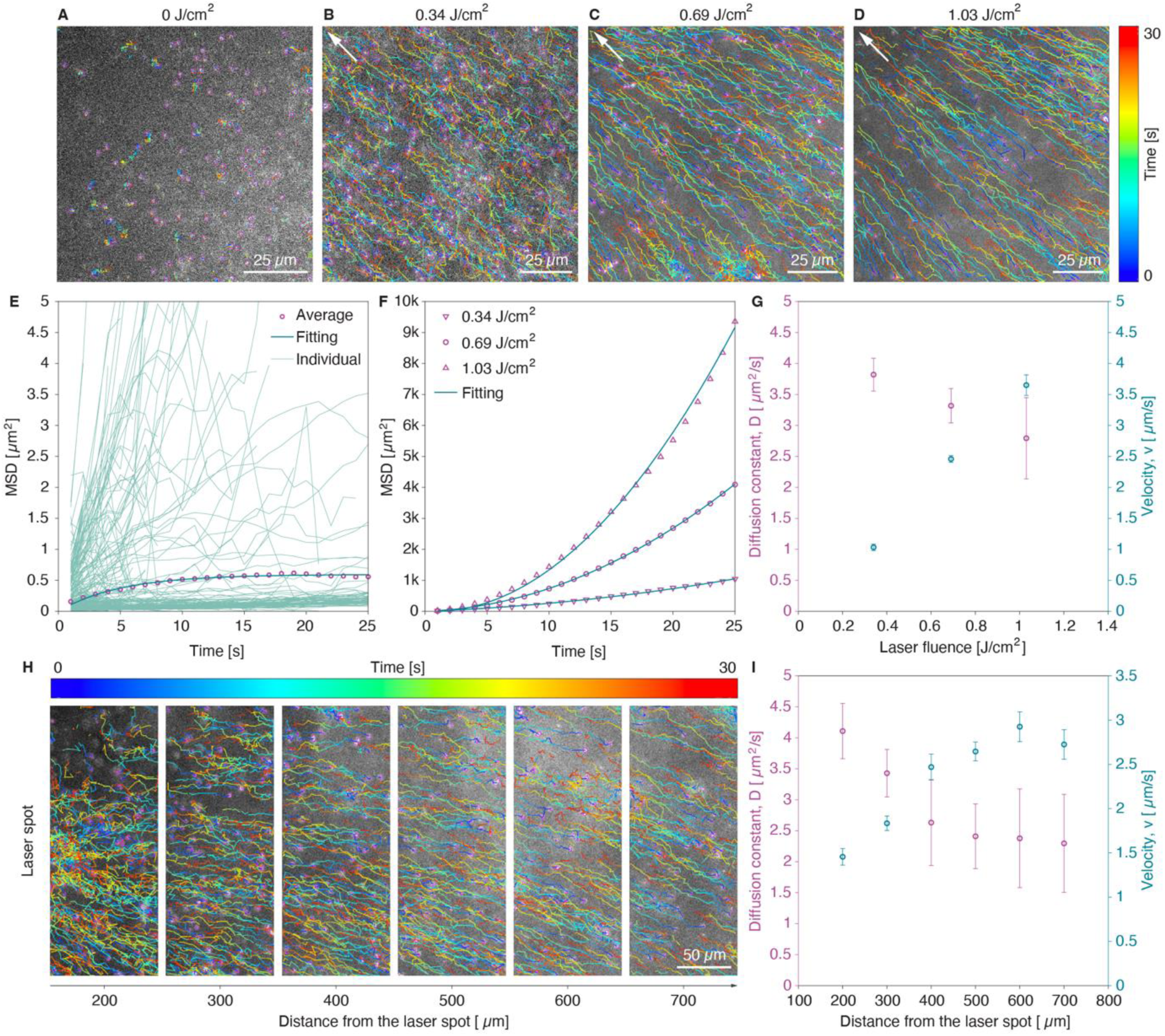
Photothermal transport of PS nanoparticles (dp = 520 nm) with ICG upon laser irradiation in bovine vitreous. **A:** Representation of colored single traces of PS nanoparticles (dilution ratio 1/1000 (v/v)) from MPT analysis of **Video S6** in vitreous samples in the presence of ICG at 0.5 mg/mL and without laser irradiation. **B-D:** Representation of colored single traces of PS nanoparticles (dilution ratio 1/1000 (v/v)) in vitreous samples in the presence of ICG at 0.5 mg/mL and with laser irradiation (laser spots are outside of the selected area of interest indicated with white arrows in the top left corner). MPT analysis has been performed on videos recorded at different laser fluences: 0.34, 0.69 (**Video S5**) and 1.03 J/cm^2^. **E:** Evolution of MSD values with time from MPT analysis of **Video S6.** Representation of individual particles traces and exponential fitting model (i.e., confined motion). Number of traces for analysis = 447. **F:** Time traces of average MSD acquired for different laser fluences extracted from the individual traces summarized in **Supplementary Figure S4**. **G:** Evolution of the obtained fitting parameters, including diffusion constant *D* (left Y axis) and particles velocity *v* (right Y axis) as a function of laser fluence. **H:** Representative colored trajectories of PS nanoparticles (dilution ratio 1:1000 (v/v)) located at distances ranging from 100 to 800 µm from the laser spot from MPT analysis of **Video S8** in vitreous samples in the presence of ICG at 0.5 mg/mL. **I:** The fitting parameters, diffusion coefficient *D* (left Y axis) and particle velocity *v* (right Y axis), plotted as a function of distance from the laser spot extracted from the MSD traces exemplified in **Supplementary Figure S5**.

**Table 1.** Influence of the laser fluence on the photothermal transport of PS nanoparticles (i.e., diffusion constant and particle velocity) in bovine vitreous humor.

| Laser fluence (J/cm<sup>2</sup>)<sup>a</sup> | Diffusion constant $D$ (μm<sup>2</sup>/s)<sup>b</sup> | Particle velocity $v$ (μm/s)<sup>c</sup> |
| --- | --- | --- |
| 0 | 0.0294 ± 0.006 | –<sup>d</sup> |
| 0.34 | 3.82 ± 0.266 | 1.03 ± 0.052 |
| 0.69 | 3.32 ± 0.277 | 2.46 ± 0.054 |
| 1.03 | 2.79 ± 0.656 | 3.65 ± 0.165 |

<sup>a</sup>determined from the formula: $Laser\ fluence\ (J.cm^{-2}) = (Energy\ at\ the\ sample\ E_{sample}\ (J)) / (\pi \times (laser\ beam\ diameter\ (cm)/2)^2)$ where $E_{sample}$ = Delivered energy (J) x conversion factor, with a laser beam diameter of 217.4 μm and a conversion factor of 2.56; <sup>b</sup>obtained from the **equation (6)** for 0 J/cm<sup>2</sup> and **equation (5)** for 0.34, 0.69 and 1.03 J/cm<sup>2</sup>; <sup>c</sup>calculated from the **equation (5)**; <sup>d</sup>no significant velocity has been observed without laser irradiation.

Next, we examined the photothermal transport of PS nanoparticles in the vitreous as a function of their distance from the laser spot. For this analysis, 520 nm-PS nanoparticles, diluted at a ratio of 1:1000 (v/v) in an aqueous solution of ICG (0.5 mg/mL), were injected into excised vitreous samples and irradiated at a laser fluence of 0.69 J/cm². After video acquisition (**Video S8**), multi-particle tracking analysis was performed in 100 µm wide regions, located 200 to 800 µm from the laser focal point, as shown in Figure 3H. The corresponding MSD traces are shown in **Supplementary Figure S5** whereas the extracted fitting parameters for each region, including diffusion constants and particle velocities, are summarized in Figure 3I. Notably, the diffusion coefficient of the PS nanoparticles in vitreous retards significantly from 4.1 ± 0.45 µm²/s to 2.6 ± 0.69 µm²/s between 200 and 400 µm from the laser spot, after which it plateaus even beyond 800 µm. This trend suggests reduced collagen network density and/or loss of structural integrity closer to the laser spot, likely due to localized heating that causes partial degradation of the surrounding collagen matrix. MSD analysis supports this observation, showing steeper slopes for particles within 100-400 µm of the laser (**Supplementary Figure S5A-C**), indicating enhanced mobility. It is interesting to note that particle velocities, however, decrease as they approach the laser spot, dropping from 2.72 ± 0.17 µm/s at 700 µm to 1.46 ± 0.09 µm/s at 200 µm (Figure 3I). Overall, the obtained results indicate a substantial temperature rise due to ICG absorption in vitreous, resulting in enhanced particle diffusion at the laser focal point and driving nanoparticle transport over long distances (up to nearly one millimeter) from the heating site.

To assess the impact of the diameter of the particles on their photothermal transport in vitreous, we performed additional experiments using 1-µm PS particles. As vitreous has a reported average mesh size of 510-565 nm,^9,11^ particles exceeding this size become immobilized within the network, ^9,11^ raising the question of whether photothermal effects would suffice to induce their motion. 1-µm PS particles were dissolved at a 1:1000 ratio (v/v) in aqueous solutions of ICG at 0.5 mg/mL before being injected into excised vitreous samples. Selected areas within the excised vitreous samples were irradiated with the pulsed laser (0.69 J/cm^2^, 532 nm) and videos were recorded (1 fps, 30 to 60 s) before, during and after laser irradiation (see **Supplementary Table 1**). Similar to smaller particles (520 nm), these larger particles were also attracted toward the laser focal point and accumulated at the illuminated zone upon laser irradiation in the presence of ICG (Figure 4A). MSD calculations obtained from MPT analysis of **Video S9** confirmed a quadratic motion of 1-µm PS particles, relatively similar to the one observed with 520-nm PS particles as shown in Figure 4B. However, the trajectory of these particles appears slightly irregular - compared to 520-nm particles - since they follow different paths throughout the vitreous, as shown by colored single particle traces in Figure 4A in comparison with elongated and straight trajectories obtained under the same conditions (i.e., see Figure 3C, 0.69 J/cm^2^, ICG at 0.5 mg/mL, and dilution ratio of particles 1/1000 (v/v)). Upon laser irradiation (0.69 J/cm^2^, 532 nm), the motion of these bigger particles has been characterized by lower diffusion constant and similar velocity than the ones measured for 520-nm particles (i.e., *D* = 2.05 ± 0.58 *vs.* 3.32 ± 0.277 µm^2^/s and *v* = 1.9 ± 0.11 *vs.* 2.46 ± 0.054 µm/s, respectively, Figure 4C), confirming the difficulty for larger particles to go through the network due to larger hydrodynamic drag.

**Figure 4.**
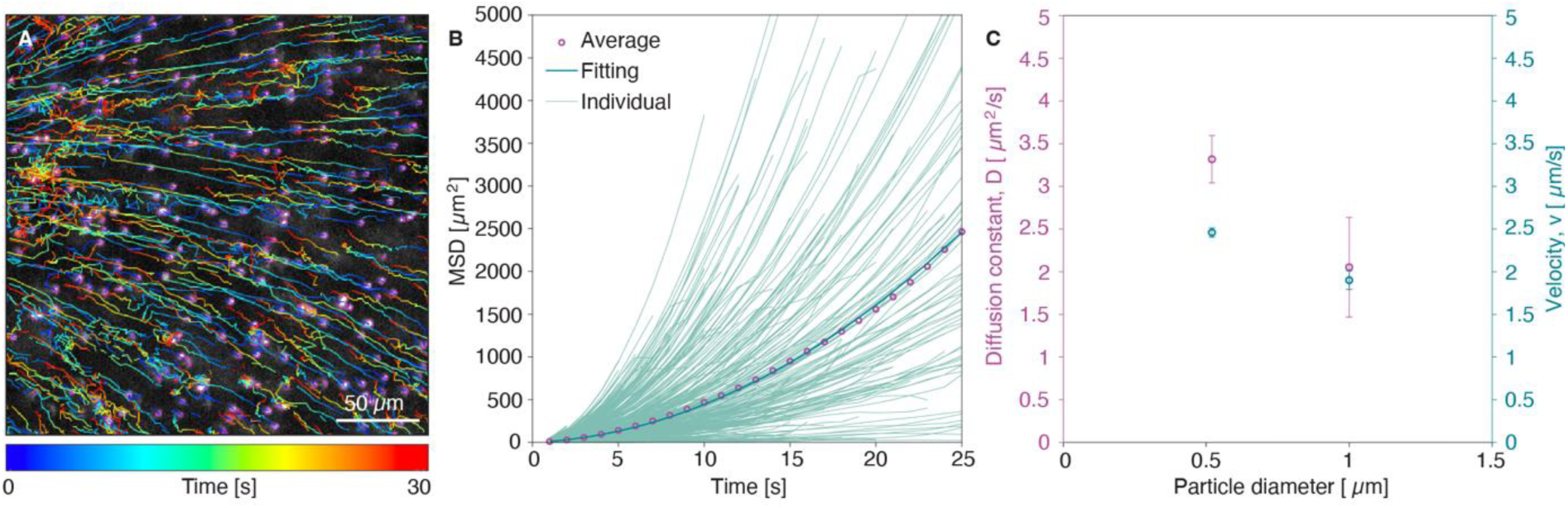
Photothermal transport of PS particles (dp = 1 µm) in the presence of ICG upon laser irradiation in bovine vitreous. **A:** Representation of colored single traces of PS particles (dilution ratio 1/1000 (v/v)) from MPT analysis of **Video S9** in vitreous samples in the presence of ICG at 0.5 mg/mL upon laser irradiation (0.69 J/cm^2^, 532 nm). The laser spot is outside of the selected area of interest in the left hand side. **B:** Evolution of MSD values with time from MPT analysis of **Video S9** with individual particle traces and quadratic fitting model. Number of traces for analysis = 926. **C:** The obtained fitting parameters, including diffusion constant *D* (left Y axis) and particle velocity *v* (right Y axis), as a function of the particle diameter.

### Numerical simulations of the photothermal effects

To investigate the underlying mechanisms driving nanoparticle motion in water and vitreous, we simulated different photothermal effects using finite-element modeling^38–41^ (see **Methods**). These simulations account for heat generation resulting from ICG light absorption under laser irradiation, and compute the resulting temperature gradients, fluid dynamics (thermal convection), and thermophoretic particle transport. We modeled the propagation of a Gaussian laser beam in paraxial approximation^42,43^ (see **Methods**) through media with a refractive index *n* = 1.33, including a 200 µm-thick water layer and a 500 µm-thick vitreous layer (Figure 5), closely matching both experimental conditions. The ICG-containing medium was modeled as a dielectric slab with absorbing properties. Upon laser illumination, this absorption leads to resistive power dissipation, resulting in a transient temperature distribution within the illuminated region (Figure 5A). To capture fluid motion induced by local heating, the convection heat transfer and incompressible Navier–Stokes equations were solved, yielding flow velocity fields corresponding to thermally driven convection (Figure 5B). In addition to thermal convection, we also calculate the contribution of thermophoresis independently, using the modelled temperature gradients ∇*T.* To compute thermophoretic velocity *v_tp_* of nanoparticles (Figure 5C) we use the following relation^21,23,27^:

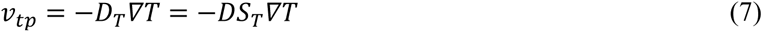

where *D_T_* is the thermophoretic diffusion coefficient, *D* is the Brownian diffusion coefficient, and *S_T_* = *D_T_* / *D* is the Soret coefficient. The Soret coefficient quantifies the strength and direction of particle movement in response to a temperature gradient and depends on many factors, including temperature, particle size, surface properties, solvent viscosity and thermal capacitance.^23,27^ In our simulations, the Soret coefficient has a negative value (see **Methods**), indicating thermophilic thermophoresis (i.e., particle migration from cooler regions toward the localized heat source).

**Figure 5.**
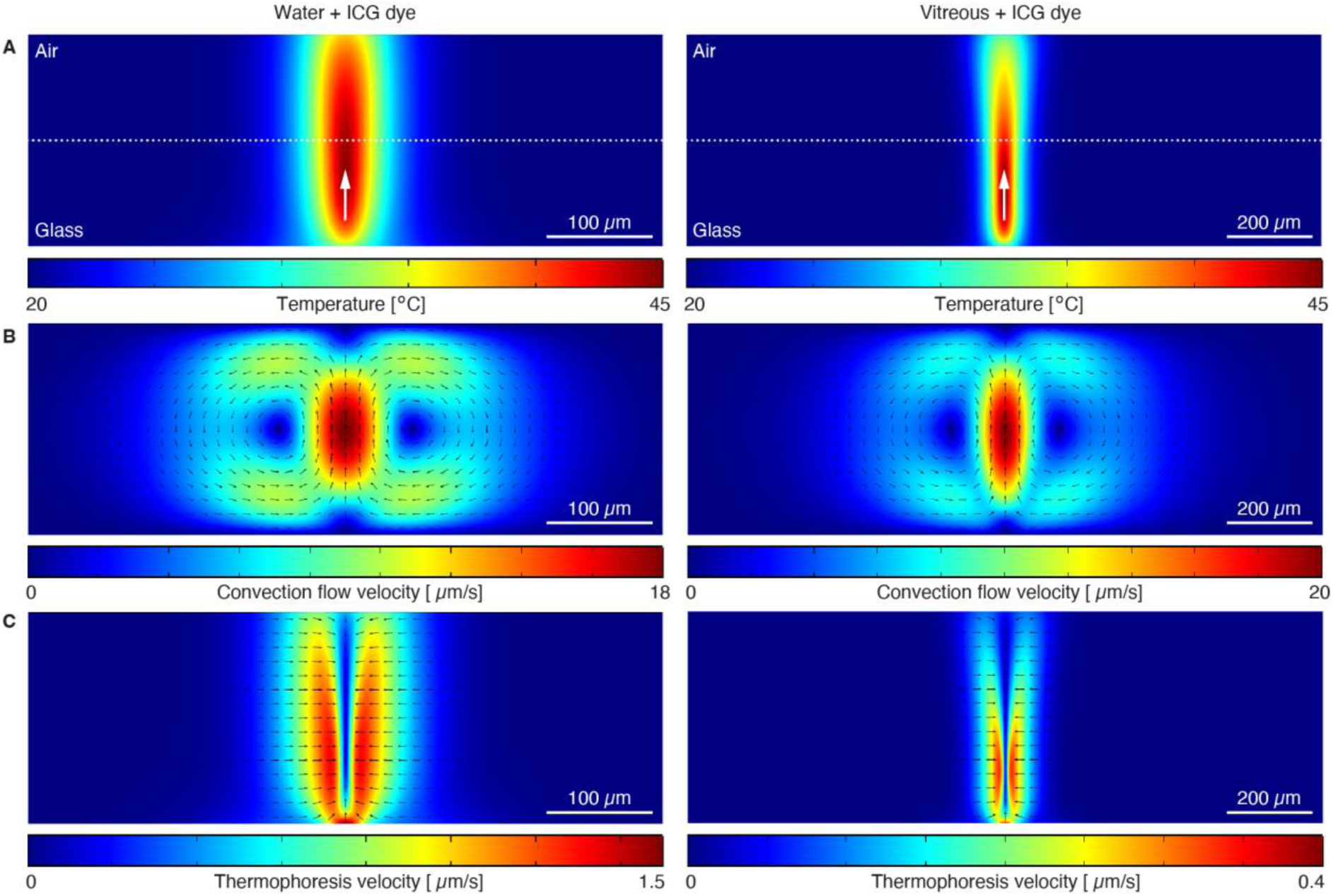
Numerical calculations of different photothermal effects induced by ICG absorption of the laser energy in water (left panels) and in bovine vitreous (right panels, x,z plane). **A:** Steady-state temperature distribution following laser irradiation shows localized heating centered around the laser beam, propagating upward as indicated by white arrow and focused at the center of the sample, as marked by the dotted line. The samples are modelled between two boundaries, with air at the top surface and glass at the bottom; as a result, the temperature profiles are asymmetric along *z*-axis due to differing thermal conductivities of air and glass. **B:** Thermal convection flow fields generated by the temperature gradient, with arrows indicating the direction and circulation pattern of fluid motion. **C:** Thermophoretic velocities of particles due to opto-thermal gradients (thermophoresis), with arrows indicating the particle motion towards the laser spot.

The calculated results reveal strong contrasts between water and vitreous environments. *In water*, the laser-induced heating produces a broad and intense temperature gradient (Figure 5A, left), leading to pronounced thermally-induced convection flows (Figure 5B, left) with recirculating patterns and peak flow velocities up to ∼18 µm/s in the hot spot. These flows, coupled with thermophoretic forces which can speed up the particles up to 1.5 µm/s (Figure 5C, left), effectively drive nanoparticles toward the laser focus, in good agreement with the experimentally observed directed motion. Similar effect is observed in *the vitreous* with localized temperature increase (Figure 5A, right) and more significant convective motion (Figure 5B, right) due to thicker sample height. Several studies have reported that thicker samples have stronger convection flows under thermal gradients which become dominant over thermophoretic motion.^19,23,44,45^ The calculated thermophoretic velocities in vitreous are reduced (Figure 5C, right), reaching only ∼0.35 µm/s compared to 1.5 µm/s in water, caused by the higher liquid viscosity. Another notable feature is that the radius of particle attraction driven by thermal convection toward the heating spot scales with the sample thickness, with flow velocities attenuating over distances of approximately 200 µm in water and 500 µm in vitreous (**Supplementary Figure S6**), whereas thermophoretic forces act over much shorter ranges, typically less than 100 µm around the laser beam (**Supplementary Figure S6**). All these findings suggest that while both convection and thermophoresis contribute to particle motion in water upon laser irradiation, particle transport in vitreous is predominantly governed by convective fluid flow with a small thermophoretic contribution.

Beyond the effects of temperature and viscosity, fluid velocity is also strongly influenced by spatial position relative to the heat source and nearby boundaries. As previously shown in Figure 5B, velocity fields vary with both the radial distance from the laser center and the vertical distance from the glass substrate. **Supplementary Figure S6** further illustrates this behavior, where extracted velocity profiles for water and vitreous reveal that particles closer to the glass surface (*z* = 10 µm) experience reduced convection velocities due to stronger viscous drag at the no-slip boundary, whereas particles further away from the surface (*z* > 20 µm) move faster. This spatial dependence explains the variations in particle velocities observed experimentally in water (Figure 2H), where speeds fluctuate around 1 µm/s for ICG concentrations between 0.25 and 0.5 mg/mL. These differences likely result from variations in the focal plane position relative to the glass surface during microscopy, as well as the distance between the analyzed region and the laser focal spot (approximately 100–200 µm), as depicted in **Supplementary Figure S6**.

### Photothermal transport of PS nanoparticles in aging vitreous (Figure 6A)

Firstly, videos were recorded from vitreous samples aged 0, 4 (**Video S10**) and 7 days (**Video S11**) containing 520 nm-PS nanoparticles (dilution ratio of 1/1000 (v/v)), but without adding ICG or laser irradiation (see **Supplementary Table 1**). These videos revealed an increased random motion of PS nanoparticles in the absence of laser irradiation, likely due to a less confined environment resulting from progressive vitreous liquefaction over time. MPT analysis (Figure 6B) yielded diffusion constants which increase with time for samples aged 0, 4 and 7 days (0.029 ± 0.006, 0.038 ± 0.003, and 0.17 ± 0.005 µm^2^/s, respectively, Figure 6C). The determined confinement radius (see **Methods**) supports the presence of a degraded microenvironment in week-old vitreous samples, exhibiting a radius of 4.5 ± 0.8 µm - approximately six times larger than that measured in freshly analyzed samples at day 0 (0.77 ± 0.06 µm). Previous studies have reported an average mesh size of 510-565 nm for native vitreous that remains intact within the eye.^9,10^ In our case, the observed higher radius values may result from mechanical deformation or structural relaxation of the vitreous matrix, related to liquefaction over time and extraction from the eye. Additionally, the slight negative surface charge of the particles may provide them with some extra mobility due to electrostatic repulsion with the vitreous matrix.^11^

**Figure 6.**
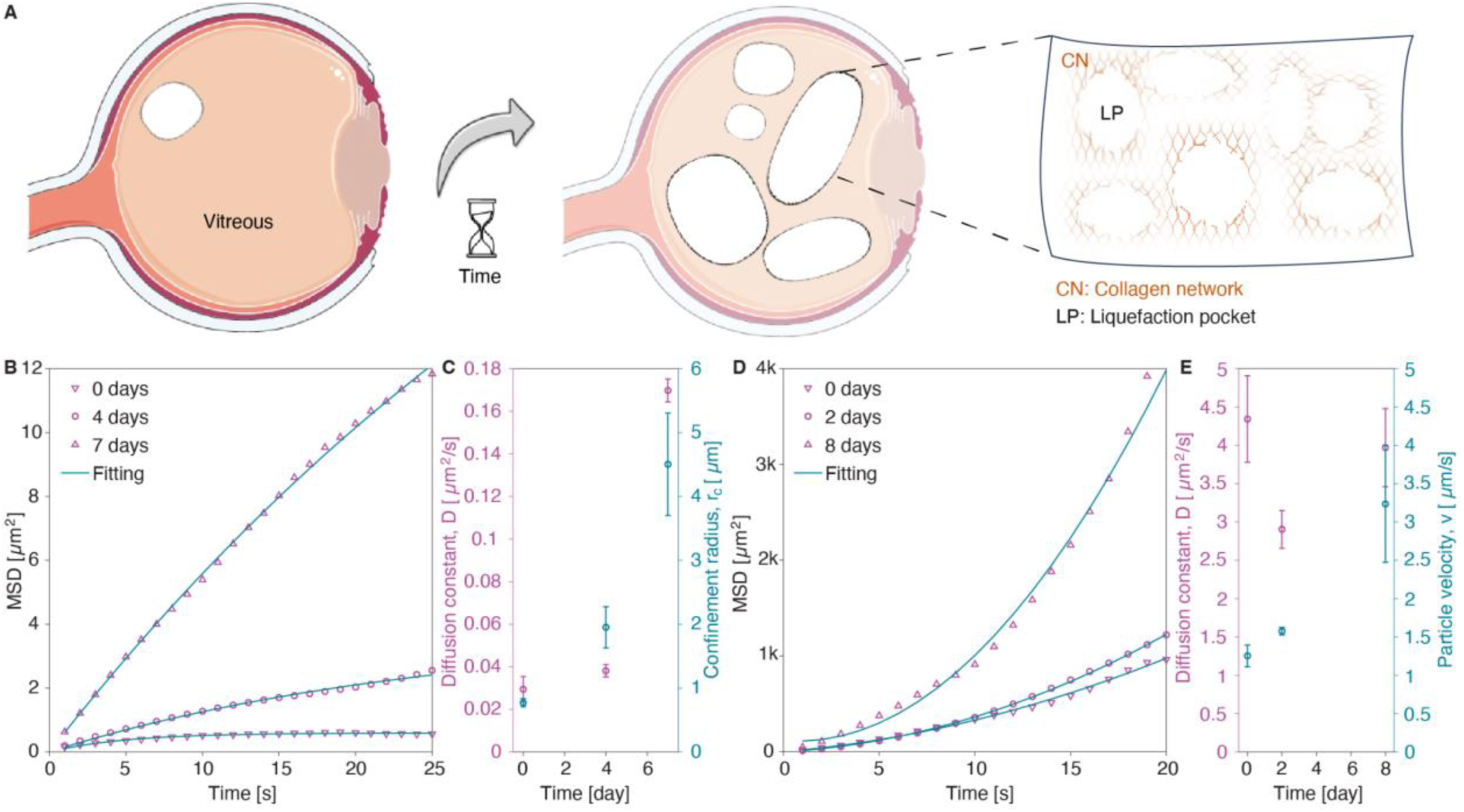
Influence of vitreous liquefaction on the photothermal transport of PS nanoparticles with and without laser irradiation. **A**: Schematic illustration of vitreous liquefaction over time (i.e., formation of liquefaction pockets (LP) and related collagen network (CN) degradation; **B**: Time traces of average MSD for aged-vitreous samples (0, 4, and 7 days post-dissection) containing 520 nm-PS particles (dilution ratio 1/1000 (v/v)) without laser irradiation or ICG, based on individual particle traces summarized in **Supplementary Figure S7**. **C**: Evolution of the obtained fitting parameters, including diffusion coefficient *D* (left Y axis) and confinement radius *rc* (right Y axis), as a function of time after vitreous extraction. **D**: Time traces of average MSD for laser-irradiated (0.69 J/cm^2^, 532 nm) aged-vitreous samples (0, 2, and 8 days post-dissection) containing 520 nm-PS particles (dilution ratio 1/1000 (v/v)) and ICG (0.5 mg/mL), based on individual particle traces shown in **Supplementary Figure S8**. **E**: Extracted fitting parameters - diffusion coefficient *D* (left Y-axis) and particle velocity *v* (right Y-axis) - as a function of time after vitreous extraction under laser irradiation.

Secondly, videos have been recorded from vitreous samples aged 0, 2 (**Video S12**) and 8 days (**Video S13**) containing 520 nm-PS nanoparticles (dilution ratio of 1/1000 (v/v)), in the presence of ICG (0.5 mg/mL) and upon laser irradiation (0.69 J/cm^2^, 532 nm) to study the influence of vitreous aging on the photothermal transport of PS nanoparticles (see **Supplementary Table 1**). The results are summarized in Figure 6D, where the average MSD curves are plotted as a function of time. The MPT analysis revealed that upon laser exposure, particles in 8 day old vitreous samples (**Video S13**) follow a quadratic motion with a significant higher velocity (3.24 ± 0.76 µm/s, see Figure 6E) than the ones determined on day 0 or 2 (**Video S12**), being 1.25 ± 0.14 µm/s and 1.58 ± 0.05 µm/s, respectively (see Figure 6E), while the diffusion constant remains similar in all samples (3.97 ± 0.51, 2.9 ± 0.25, 4.34 ± 0.56 µm^2^/s for 8-day, 2-day, and 0-day samples, respectively) as shown in Figure 6E.

These two experiments (i.e., without and with laser irradiation) demonstrate a significant effect of vitreous aging on the mobility of nanoparticles without laser irradiation (i.e., by an increase of the confinement radius) and with laser irradiation (i.e., by an increase of the particle’s directional velocity), consistent with progressive vitreous liquefaction. It proves that vitreous liquefaction is a non-negligible factor in the photothermal transport of nanoparticles in the vitreous. Importantly, however, liquefaction did not impede the guidance of particles under laser irradiation.

## Discussion

In this study we have demonstrated that the combined use of a light absorbing ocular dye (ICG) and a nanosecond pulsed-laser irradiation (2-5 ns) allows photothermal transport of PS nanoparticles in both water and bovine vitreous. The particles exhibit directed motion toward the laser focus, forming elongated trajectories, whereas in the absence of laser irradiation and/or ICG, their motion remains Brownian (random) and even more restricted in the tight matrix of the vitreous (confined motion). We observed particle accumulation (for both the 520-nm and 1-µm PS particles) at the laser focus point in both media. Varying experimental parameters such as ICG concentration, laser fluence, particle diameter and distance from the laser focus point, also impacted the photothermal transport of the particles. The particle velocity can be therefore adjusted by changing the dye concentration. We identified that a minimal concentration of ICG (0.2 mg/mL) is needed to induce photothermal transport; at lower ICG concentrations the PS nanoparticles only exhibited Brownian motion while upon increasing the ICG concentration, transport of PS nanoparticles, through both thermal convection and thermophoresis, became stronger, highlighting the crucial role of ICG. The laser fluence also plays a crucial role in the photothermal transport of nanoparticles as a monotonic increase in particle velocity with fluence has been observed. While the mesh size of the collagen network in the vitreous is around 500 nm, the motion of larger particles (1 µm) seemed possible upon pulsed-laser illumination in the presence of ICG. We also want to raise attention on the importance of the particle distance from the laser spot in photothermal transport. Beyond the successful transport of nanoparticles over long distances (up to nearly one millimeter) from the heating site, we observed enhanced particle diffusion near the laser focal point, resulting from localized heating and partial disruption of the vitreous network. We also demonstrated that the velocity tends to decrease as particles approach the laser focus, likely due to a high accumulation which restricts mobility in the progressively crowded environment. This observation suggests that excessive accumulation near the illuminated zone creates a local crowding that hinders further motion. Our analyses of the underlying parameters influencing photothermal transport of nanoparticles might form the basis for the fine tuning of NP motion in vitreous in future medical applications. Furthermore we observed that photothermal transport of nanoparticles in the vitreous is influenced by the aging (liquefaction) of the vitreous over time, indicating that patient-to-patient variability (age, pathology, etc.) might impact the photothermal transport of particles.

Building on our empirical findings we developed a numerical model that offers valuable insights into the mechanisms underlying photothermal transport. We found that the photothermal transport of nanoparticles in both water and vitreous is governed by a combination of thermal convection and thermophoresis, with the relative contribution of each mechanism depending on the medium. In the vitreous, the thermophoretic velocity is markedly reduced due to its higher viscosity, while stronger convection flows emerge as a result of the greater sample thickness compared to water. Importantly, fluid velocity also strongly depends on the spatial position relative to the heat source and nearby boundaries. Although simulated velocities near the glass surface are lower in the vitreous than in water (**Supplementary Figure S6**), experimental data reveal higher particle velocities in the vitreous (Figure 3G). This apparent discrepancy can be explained by the increased sample thickness in vitreous (exceeding 500 µm), which supports more extensive convection flows and a longer-range particle attraction toward the laser spot. For example, in Figure 3H, particles are drawn in from distances beyond 800 µm, suggesting a substantial vertical extent of the sample. Additionally, prolonged laser exposure (over 30 s) can lead to cumulative heating, resulting in significant local temperature increases and partial liquefaction of the vitreous, thereby reducing its viscosity and further enhancing fluid motion. Together, these theoretical results highlight the complex interplay of thermal convection, thermophoresis, and boundary conditions, all of which must be carefully considered when interpreting particle transport under opto-thermal excitation, particularly for biomedical applications.

Our experimental and numerical findings prove to be promising for light-mediated delivery in the posterior part of the eye but further work is needed for future clinical implementation, especially regarding laser use and heat generation. In general, heat generation under laser irradiation is a cumulative process; over prolonged exposure, local temperatures progressively rise, leading to steeper thermal gradients and stronger fluid flows. In our simulations, the temperature increase shown in Figure 5A was calculated over a 10 ms period, corresponding to the 100 Hz laser repetition rate. When we extend the simulations to longer timescales, such as 0.1-1 s, the cumulative heating can elevate local temperatures to nearly 100 °C, potentially leading to liquid boiling. However, in our experiments, boiling was rarely observed (only for very high laser fluences) because continuous fluid motion brings cooler liquid toward the laser spot, mitigating local temperature rise. Simulated heating can also be adjusted by varying the absorption coefficient (ICG concentration) or the laser power (fluence). This aligns with experimental results, where particle diffusion constants in water increase with higher ICG concentrations (Figure 2H), corresponding to local temperatures approaching 80 °C at 0.5 mg/mL, or with increasing laser fluence (Figure 3G). In our study, a pulsed-laser (2-5 ns; 532 nm) was used. A strong advantage of using this type of laser for ophthalmological purposes is that it is widely used in the clinic^46^ and that our technology would not need extensive development of optics in case of clinical translation. However, safety concerns still need to be addressed, especially concerning potential toxicity coming from the dye. Our model dye, ICG, has been approved for medical use by the Food and Drug Administration (FDA) since 1959. While our choice for this conceptual study relied on the clinical use of ICG, it is also known to induce radical oxygen species (ROS) upon light irradiation^47^ which raises potential safety concerns, particularly to the retina. However, the lack of a consistent trend in experimental results on ICG toxicity^48–50^ makes it challenging to draw any definitive general conclusion on the subject. Anyhow, to mitigate the toxicity risk of ICG, general safety guidelines have emerged like reducing the dye concentration and the exposure time.^51^ Reducing ICG concentration from 5 to 0.5 mg/mL^-1^ and the exposure time (less than 10 s) enabled surgeons to suppress visual field defects and retinal pigment epithelium (RPE) damages for inner limiting membrane (ILM) visualization.^52^ We have demonstrated, in our approach, that optimal photothermal transport of nanoparticles can be reached for ICG concentrations starting at 0.2 mg/mL and using a pulse laser irradiation between 2 and 5 ns. In addition, another strategy to reduce potential ICG toxicity relies on increasing the osmolarity of the particle solution and/ or on modifying the dye composition. For example, it has been shown that using infracyanine green— a dye that is structurally close to ICG— dissolved in glucose 5 (w/v)% as an iso-osmolar solution, significantly reduced toxicity to RPE cells.^53^ Finally, laser irradiation can also be associated with potential toxicity risks related to the wavelength and exposure time, that are important to consider for potential clinical translation. Our study has been performed at 532 nm because this wavelength is commonly used with nanosecond lasers in the clinic (e.g. for iridotomy,^46^ laser trabeculoplasty (SLT)^54^). It is important to underline that 532 nm is not the maximum absorption peak of ICG (i.e., 780 nm^55^). Therefore, the laser irradiation of a mixture of ICG and PS nanoparticles at higher wavelengths could further stimulate the motion (i.e., diffusion and velocity) but could also lead to increased heat dissipation and potentially higher toxicity, since we have shown that laser heating increased linearly with the laser wavelength (**Supplementary Figure S9C**). Important to note as well, is that the use of 532 nm pulsed-laser in the vitreous poses risks for the retina owing to its pigmented (hence light-absorbing) nature. Some reports have indeed demonstrated that pulsed-laser irradiation in the direct neighborhood of the retina could induce photomechanical damage.^56^ Therefore, other dyes with limited ROS generation and absorption in the near infrared (NIR) II region could be considered to further improve the safety of our approach by limiting photomechanical damage. Finally, the laser fluence also plays an important role for safety. The fluence range used in this study falls within the same order of magnitude as the fluences typically applied in clinical settings with 532 nm-YAG lasers (e.g., Ellex Tango Reflex), which range from approximately 0.2 to 2 J/cm².^57^ We demonstrated that fluence values within this range (i.e., 0.69 and 1.03 J/cm², see **Table 1**) are already sufficient to induce photothermal transport of PS nanoparticles in the presence of ICG in vitreous. This suggests that effective photothermal transport can be achieved at clinically relevant fluences, potentially allowing for safe translation. However, further research is yet still needed to identify the optimal balance between safety and efficacy (i.e. reaching high velocity at minimal energy).

Given its significant physical and experimental insights, we believe this first study on the ‘directed’ photothermal transport of particles in the vitreous, using clinical laser settings and a clinically approved dye, might pave the way for light-guided strategies to deliver intravitreally-injected drugs to the posterior segment of the eye. Our approach offers interesting therapeutic perspectives to face the major challenges associated with the intravitreal injection of drugs and particles, e.g. an insufficient drug concentration in target tissues caused by random and non-directed diffusion. By enabling more precise drug delivery, our strategy could ultimately reduce the need for frequent IVT injections, such as the bi-monthly injections currently required for wet age-related macular degeneration (AMD) treatment.^58^ In doing so, it may not only improve patient compliance and therapeutic outcomes, but also significantly lower healthcare costs by reducing the number of administrations of often expensive drugs.

## Materials & Methods

### Chemical materials

Deionized water was obtained from a Milli-Q water purification system (Millipore, Bedford, MA, USA). Indocyanine green (ICG) was purchased from U.S. Pharmacopeia (USP, United States). Fluorescent Negatively-Charged Polystyrene (PS) Microspheres, Dragon Green (λ_ex_=480 nm, λ_em_=520 nm) 0.50 µm and 1.0 µm were purchased from Bangs Laboratories, Inc.

### Water sample preparation

Physical mixtures of PS nanoparticles and ICG were prepared by first diluting ICG in deionized water using vortexing (Vortex V-1 plus, BioSan) for 10-20 sec. Then, PS nanoparticles have been diluted into ICG aqueous solutions and subsequently vortexed for another 10-20 sec. The volume of PS nanoparticles has been adjusted depending on the targeted dilution ratio, e.g. for 1/1000 (v/v), 1 μL of PS nanoparticles has been added into 999 uL of ICG aqueous solution. Two droplets (around 100 μL) of PS nanoparticles-ICG aqueous mixture of ICG and PS nanoparticles were placed on a microscope slide pre-treated with glue sticker, then covered with a cover glass to minimize water evaporation during experiments prior to applying nanosecond laser pulses (Ekspla N230-100-SCU-2H OPO, 532 nm, 2-5 ns). See **Video S14** (**Supplementary Table 1**) as an example with a full field of view..

### Bovine vitreous extraction and sample preparation

Fresh bovine eyes were enucleated only a few hours after cows were slaughtered (slaughterhouse Vion Group, Zotegem, Belgium). The eyes are transported and kept in ice cold CO_2_ independent medium until dissection. After removing all extra-ocular connective tissue and disinfecting the eyes by soaking them in 20 % ethanol, the sclera is punctured with a 21G needle around 10 mm below the limbus. This hole next serves as an entry point for the scissors used to cut the eye into two parts. Since the vitreous has a very fragile structure, it was carefully removed from the anterior part using a brush and kept at 4°C before use. First, 300 - 500 μL of vitreous were carefully cut and placed on a 50 mm glass-bottomed dish. Subsequently, 50 μL of the aqueous mixture of ICG and (520 nm or 1 μm) PS nanoparticles were injected in the vitreous sample using a 1 mL syringe equipped with a 21.5G needle. The sample was allowed to equilibrate for 5 minutes with a cover slip to minimize water evaporation during experiments prior to applying nanosecond laser pulses (Ekspla N230-100-SCU-2H OPO, 532 nm, 2-5 ns). See **Video S15** (**Supplementary Table 1**) as an example with a full field of view.

### Bovine vitreous liquefaction and sample preparation

Fresh bovine eyes were enucleated only a few hours after cows were slaughtered (slaughterhouse Vion Group, Zotegem, Belgium). The eyes are transported and kept in ice cold CO_2_ independent medium until dissection. After removing all extra-ocular connective tissue and disinfecting the eyes by soaking them in 20 % ethanol, the sclera is punctured with a 21G needle around 10 mm below the limbus. This hole next serves as an entry point for the scissors used to cut the eye into two parts. Since the vitreous has a very fragile structure, it was carefully removed from the anterior part using a brush and kept at 4°C before use. 300 - 500 μL of vitreous at days 0, 2, 4 and 8 were carefully cut and placed on a 50 mm glass-bottomed dish. The sample preparation was then carried out as described previously for vitreous samples.

### Optical setup and camera recording

An inverted microscope (TE2000-E, Nikon ECLIPSE) equipped with a 10x objective (Nikon CFI Plan Fluor) was used to focus the nanosecond laser on the areas containing PS nanoparticles and ICG within the samples (**Supplementary Figure S10**). As PS nanoparticles scatter sufficient light, dark-field microscopy was performed to visualize them. Samples were then illuminated with nanosecond laser pulses at a wavelength of 532 nm with a pulse duration of 2 to 5 ns (Ekspla N230-100-SCU-2H OPO) and repetition rate of 100 Hz. A beam expander (#GBE05-A, Thorlabs) combined with iris diaphragm (#D37SZ, Thorlabs) were used to adjust the diameter of the laser beam to 203 µm. The laser pulse energy was monitored by an energy meter (J-25MB-HE&LE, Energy Max-USB/RS sensors, Coherent) synchronized with the pulsed laser and placed behind a beamsplitter (**Supplementary Figure S10**). The laser fluence has been determined based on the energy value and the laser spot size, based on the following expression: *Laser fluence (J.cm^-2^) = (Energy at the sample E_sample_ (J)) / (π × (laser beam diameter (cm)/2)^2^) where E_sample_ = Energy at the energy meter (J) × conversion factor*. The conversion factor takes into account the optical path in between the beamsplitter and the sample. The set-up was adapted to record videos before, during and after laser illumination using a sCMOS camera (Photometrics Prime sCMOS, pixel size of 6.5 µm) and Micro-Manager software (Multi-D acquisition mode). A 10x microscope objective has been used (pixel size of 6.5/10 = 0.65 µm, pixel area of 1316 µm x 1316 µm). 30 to 60 seconds videos have been recorded using a frame rate of 1 frame per seconds (fps).

### Multi-particle tracking (MPT) analysis

Particle motion was analyzed using multi-particle tracking algorithms^11,42,43,44^ applied to time-lapse microscopy videos acquired under various experimental conditions. The two-dimensional positions of PS nanoparticles were tracked frame-by-frame using Fiji extension^66^ based on ImageJ and a custom-written MATLAB script, allowing for the reconstruction of individual particle trajectories over time. The mean square displacement for each trajectory was calculated using equation (3). Depending on the observed motion type, different fitting models^47^ were applied: equation (4) was used for free Brownian motion (Figure 2E), equation (5) for directed motion (Figure 2F), and equation (6) for confined motion (Figure 3E). Ensemble-averaged MSD curves were generated from multiple trajectories under each condition, and the resulting data were fitted accordingly. Based on the fitting, the key fitting parameters were extracted, including the diffusion coefficient *D*, confinement radius *r_c_*, and particle velocity *v*. Statistical error bars were derived from the fitting residuals across individual particle tracks. To estimate the local temperature from the measured diffusion coefficients, we used equations (1) and (2) to generate a diffusion-temperature calibration curve (**Supplementary Figure S9A**), based on the temperature-dependent dynamic viscosity of water.^67^ Using this curve, the diffusion constants extracted from experimental data (Figure 2F) were mapped to their corresponding temperatures, as shown in **Supplementary Figure S9B**.

### Numerical calculations of photothermal effects

Numerical simulations were performed using finite-element solver COMSOL Multiphysics 5.6 according to our previously published methods.^45,46,47,48^ To investigate the photothermal effects induced by laser-illuminated ICG in different media, including water and bovine vitreous, the simulation model coupled several different modules, including electromagnetic waves (EMW), heat transfer (HT) and computational fluid dynamics (CFD), to capture the relevant mechanisms driving nanoparticle motion. The laser was modeled as a Gaussian beam in paraxial approximation^49,50^ with an average power *P* = 100 mW, corresponding to a fluence of 2 J/cm² over a 270 µm-diameter spot (area ≈ 5.73×10^-8^ m²), yielding energy ∼1.15 mJ per pulse at repetition rate *f* = 100 Hz. This results in an average power of ∼115 mW, matching experimental conditions. To justify treating the pulsed laser as a continuous source, we computed the thermal impedance *Z = (ωC*k*)^-1/2^* for the applied materials, where ω = 2π×100 Hz is the angular frequency of the laser pulses, *C* = 4.2 MJ/(m³·K) is the volumetric heat capacity of water, and *κ* = 0.6 W/(m·K) is its thermal conductivity. This gives *Z* ≈ 2.5×10*^-5^* K/W, leading to negligible per-cycle temperature fluctuations (∼2.5 µK) relative to the total rise (∼45 °C over 10 ms), thus validating the continuous-wave approximation. The medium (refractive index *n* = 1.33) was modeled as a dielectric slab with a height *h* = 200 µm for water or 500 µm for vitreous, and a lateral width of 3*h*. The dynamic viscosity of liquefied vitreous was assumed to be four times that of water, based on reported values: 3 mPa s for vitreous^68^ and 0.7 mPa s for water^67^ at 37 °C. Laser absorption by ICG was simulated using an effective electrical conductivity σ = 0.5 S/m, chosen to match the experimental power (100 mW) and wavelength (532 nm) while ensuring that temperature rises remained below 100 °C per pulse cycle (10 ms) as shown in **Supplementary Figure S8C**. Absorption-induced heating generated spatial temperature gradients, which were then used to calculate thermal convection using the Navier–Stokes equations for incompressible flow based on the applied boundary conditions and starting room temperature of 20 °C. Thermophoretic particle motion was modeled based on these gradients using equation (7), with Soret coefficient:^32^

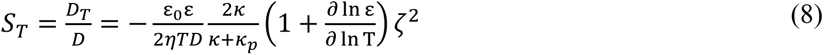

where *ε_0_* is a vacuum permittivity, *ε* = 78 is a dielectric constant of the medium, *κ_p_* = 0.13 W/(m·K) is a thermal conductivity of the particle, ζ = -30 mV is a zeta potential of the particle and τ = ∂ (ln ε) / ∂(ln T) = 2 is permittivity-related term^35^. In our case, this equation results in *S_T_* = -3.5 1/K for water and -0.8 1/K for vitreous, reflecting stronger thermophilic behavior in water due to lower viscosity. The simulations produced steady-state temperature distributions, flow velocity fields, and thermophoretic velocity profiles (Figure 5), allowing for direct comparison of the dominant transport mechanisms in each medium.

## Acknowledgment

L.G. and F.S. acknowledge the European Research Council (ERC) for funding received under the European Union’s Horizon Europe research and innovation program (Grant agreement No. 101075873, DYE-LIGHT). Y.U. acknowledges his FWO Postdoc Fellowship (1202225N).

## Data availability

All data supporting the findings of this study are available within the paper and its Supplementary Information. Source data are provided with this paper. Any further related information can be provided by the corresponding author upon reasonable request.

## Author contributions

F.S. conceived and designed the research; L.G., Y.U. and F.S. designed the experiments; L.G. and L.D.C. performed the experiments; Y.U. and L.D.C performed MPT analysis; Y.U. and F.B. developed the numerical model; L.G., Y.U. and F.S. analyzed the data and wrote the manuscript; F.S. supervised the project. All authors contributed to the discussion of the results and to the revision of the manuscript.

## Additional information

Generative AI (ChatGPT4) has been used to improve English in the manuscript. After using this tool/service, the authors reviewed and edited the content as needed and took full responsibility for the content of the publication.

