## Supplementary information for "Photothermal transport for guiding nanoparticles through the vitreous humor"

Electronic Supplementary Information for

**Supplementary Table 1. Detailed overview of the videos illustrating the photothermal transport of PS nanoparticles in both water and bovine vitreous.** Information about the medium, particles dilution ratio and diameter, dye concentration, laser fluence and aim of each experimental condition are reported in this table.

| Video (n°) | Medium | Particles | | ICG (mg/mL) | Laser fluence (J/cm$^2$) | Description (aim of the study) |
|---|---|---|---|---|---|---|
| | | Dilution ratio (v/v) | Diameter (nm) | | | |
| S1 | Water | 1/1000 | 520 | 0.5 | 2.07 | Photothermal transport in water |
| S2 | Water | 1/1000 | 520 | / | 2.07 | Influence of ICG in water |
| S3 | Water | 1/1000 | 520 | 0.5 | / | Influence of laser irradiation in water |
| S4 | Water | 1/5000 | 520 | 0.1 | 2.07 | Influence of ICG concentration in water |
| S5 | Vitreous | 1/1000 | 520 | 0.5 | 0.69 | Photothermal transport in vitreous |
| S6 | Vitreous | 1/1000 | 520 | 0.5 | / | Influence of laser irradiation in vitreous |
| S7 | Vitreous | 1/1000 | 520 | / | 0.69 | Influence of ICG in vitreous |
| S8 | Vitreous | 1/1000 | 520 | 0.5 | 0.69 | Influence of the distance from the laser spot in vitreous |
| S9 | Vitreous | 1/1000 | 1000 | 0.5 | 0.69 | Influence of the particle diameter in vitreous |
| S10 | Vitreous | 1/1000 | 520 | / | / | Influence of vitreous liquefaction without laser irradiation (4 days) |
| S11 | Vitreous | 1/1000 | 520 | / | / | Influence of vitreous liquefaction without laser irradiation (7 days) |
| S12 | Vitreous | 1/1000 | 520 | 0.5 | 0.69 | Influence of vitreous liquefaction with laser irradiation (2 days) |
| S13 | Vitreous | 1/1000 | 520 | 0.5 | 0.69 | Influence of vitreous liquefaction with laser irradiation (8 days) |
| S14 | Water | 1/1000 | 520 | 0.5 | 2.07 | Photothermal transport in water with a full field of view |
| S15 | Vitreous | 1/1000 | 520 | 0.5 | 0.69 | Photothermal transport in vitreous with a full field of view |

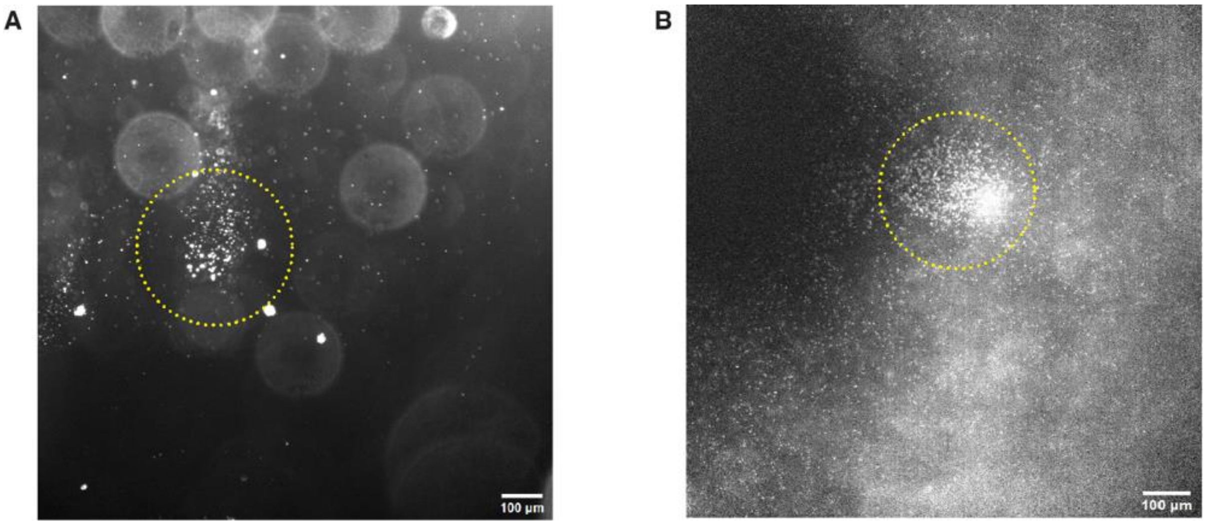


**Supplementary Figure S1. Accumulation of PS particles (1/1000 (v/v)) due to photothermal transport upon laser irradiation.** Dark field microscopy images of particle accumulation in the laser spot (indicated with yellow dotted circle) in (**A**) water (2.07 J/cm$^2$, 532 nm) and (**B**) vitreous (0.69 J/cm$^2$, 532 nm) with ICG (0.5 mg/mL).

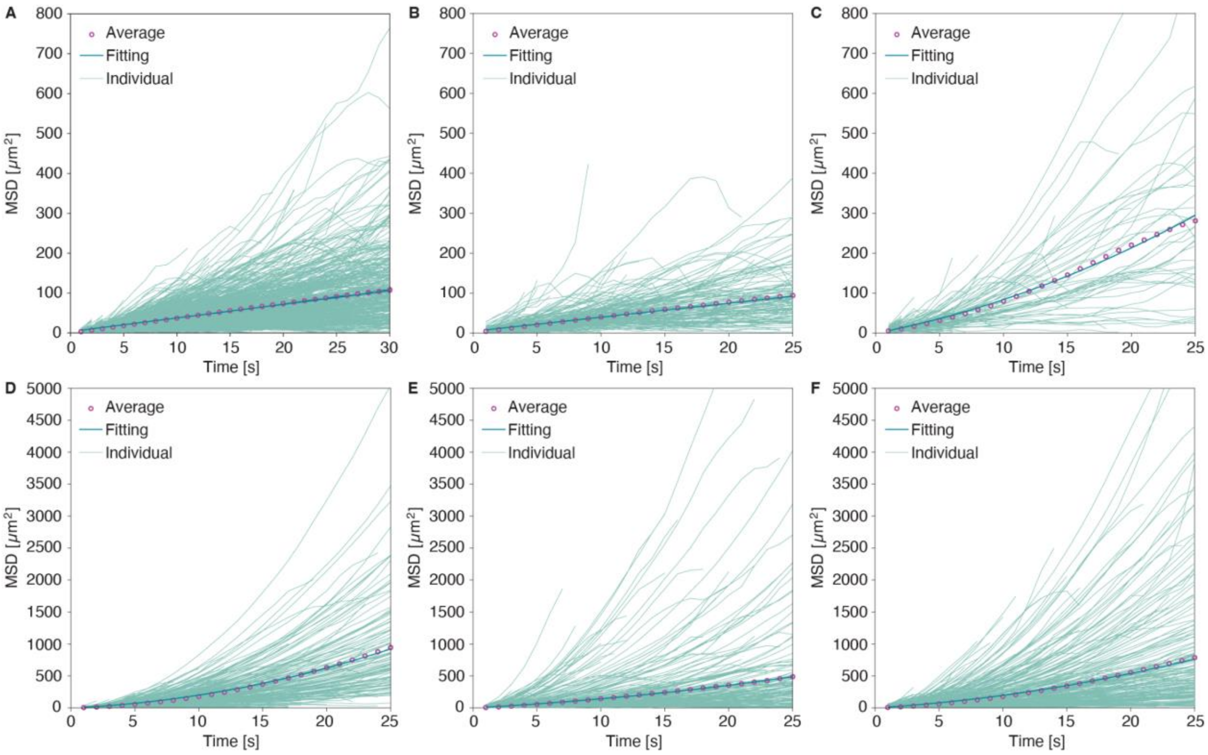


**Supplementary Figure S2. Photothermal transport of PS nanoparticles (1/5000 (v/v)) upon laser irradiation (2.07 J/cm$^2$, 532 nm) in water with varying ICG concentrations**. MSD curves as a function of time were obtained from MPT analysis for ICG concentrations of 0 (**A**), 0.1 (**B**), 0.2 (**C**), 0.25 (**D**), 0.4 (**E**), and 0.5 (**F**) mg/mL. The number of particle trajectories analyzed for each condition were: 1318, 286, 156, 760, 368, and 577, respectively.

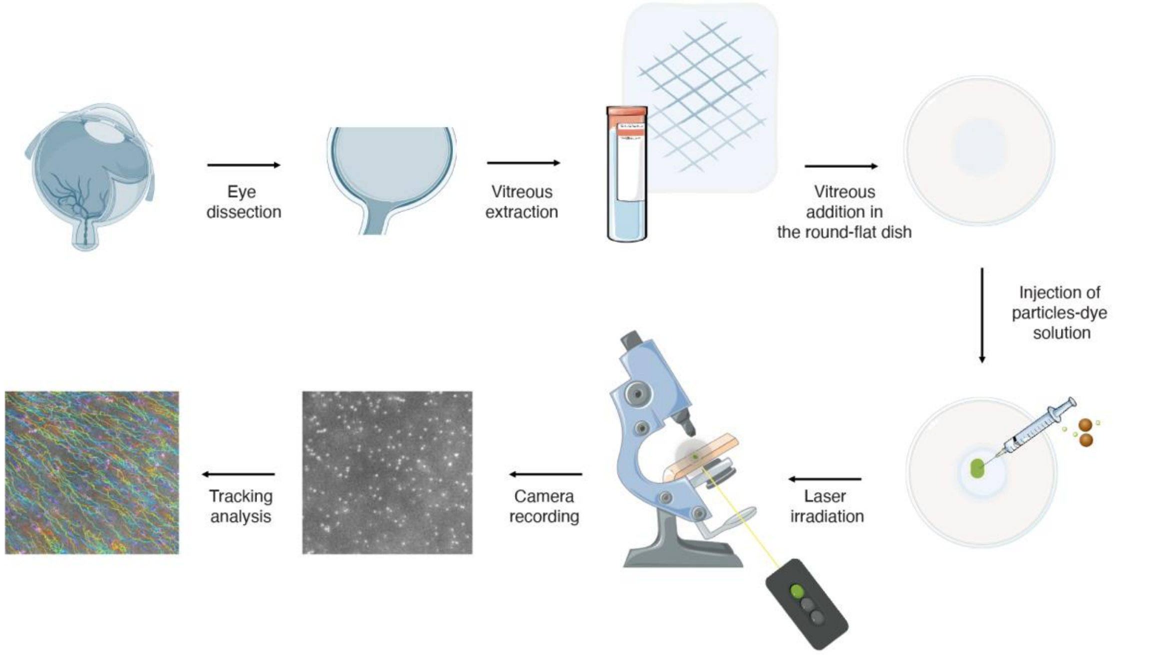


**Supplementary Figure S3. Chronological illustration of the preparation of vitreous samples for experiments.** After collection of the eyes at the slaughterhouse, the vitreous was extracted from bovine eyes and stored at 4 °C. Then, a 1 mL- droplet of vitreous was placed on a round-flat dish prior to injection of aqueous mixtures of PS nanoparticles and ICG. Finally, laser illumination was applied onto the vitreous sample and a camera was used for recording videos (before, during and after laser irradiation) and multiple-particle tracking (MPT) analysis of the videos was performed.

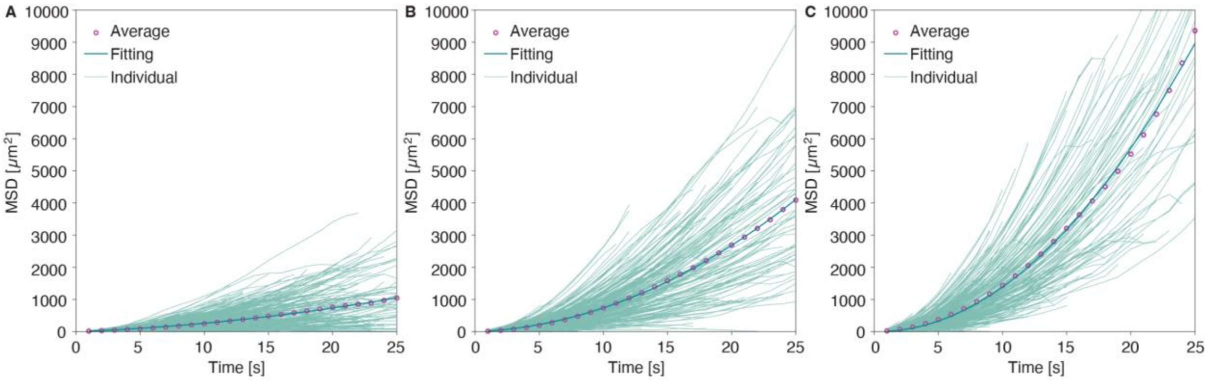


**Supplementary Figure S4. Photothermal transport of PS nanoparticles (1/1000 (v/v)) in vitreous containing ICG (0.5 mg/mL) with varying laser fluences.** Mean square displacement (MSD) curves as a function of time were obtained from multi-particle tracking analysis at three different laser fluences: 0.34 J/cm² (**A**), 0.69 J/cm² (**B**), and 1.03 J/cm² (**C**). The number of particle trajectories analyzed for each condition was 1230, 726, and 532, respectively.

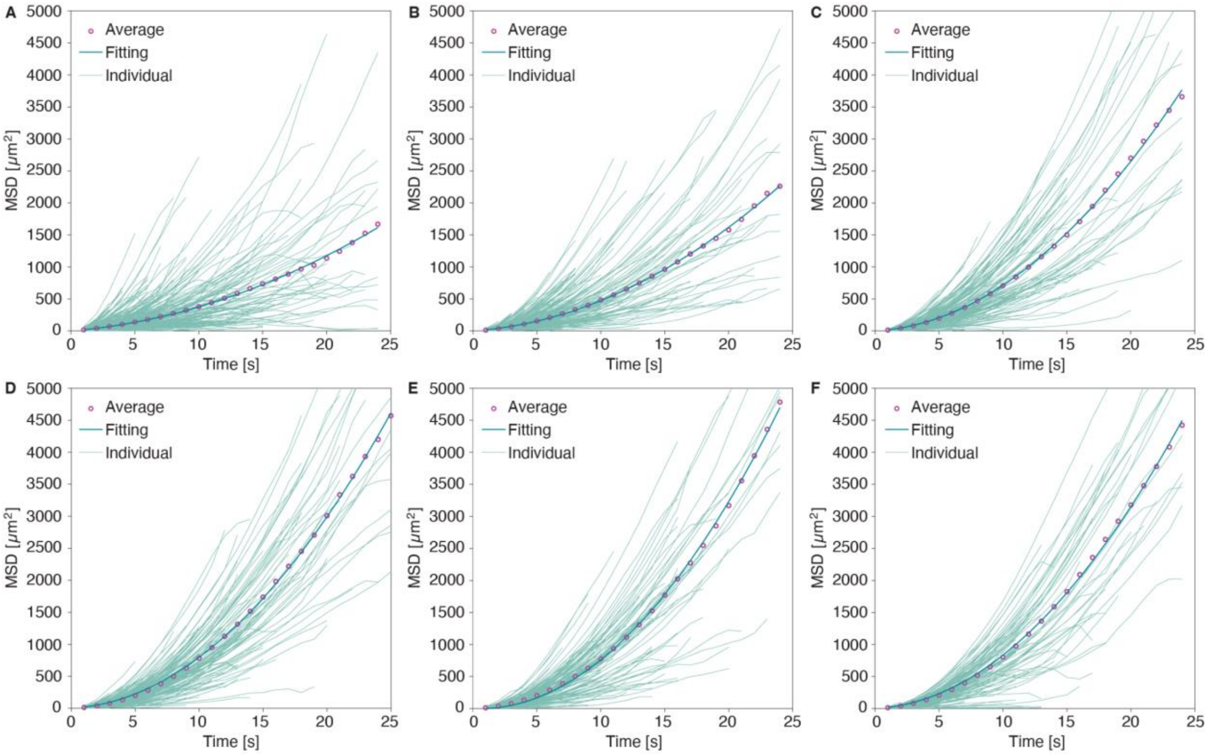


**Supplementary Figure S5. Photothermal transport of PS particles (1/1000 (v/v)) under laser irradiation (0.69 J/cm$^2$, 532 nm) in vitreous as a function of distance from the laser spot**. MSD curves as a function of time were obtained from MPT analysis for different distances of 200 µm (**A**), 300 µm (**B**), 400 µm (**C**), 500 µm (**D**), 600 µm (**E**), and 700 µm (**F**) away from the laser focal point. The number of particle trajectories analyzed for each region were: 479, 418, 311, 285, 242, and 190, respectively.

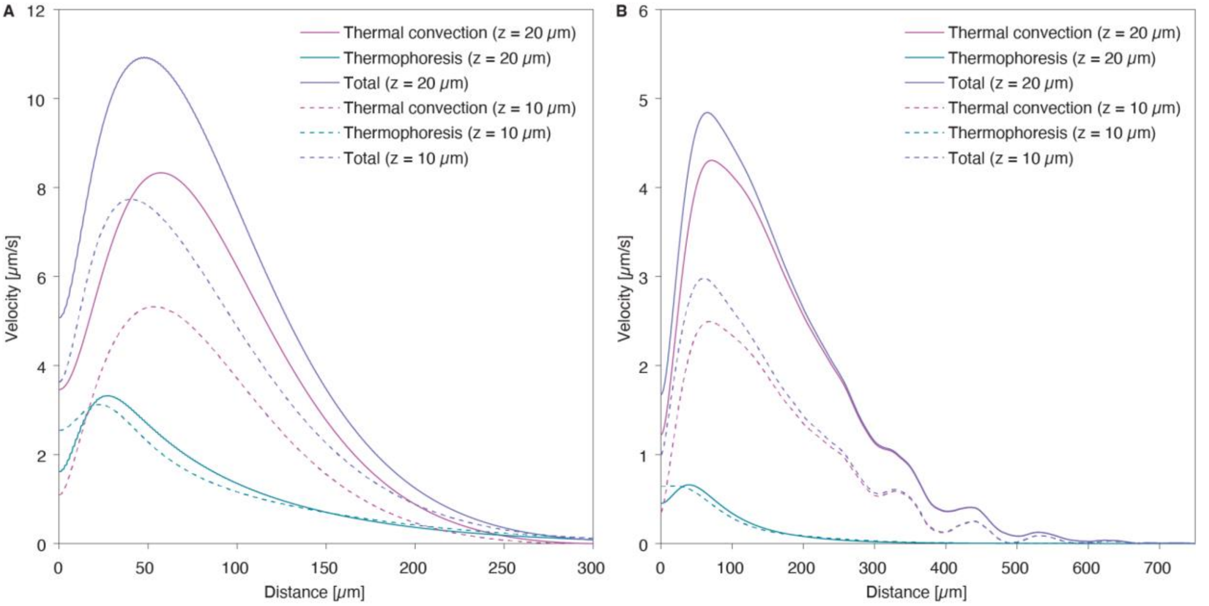


**Supplementary Figure S6. Simulated velocities of particle motion induced by thermal convection and thermophoresis as a function of distance from the laser beam center for two different heights above the surface (*z* = 10 and 20 µm). A:** Calculated velocities in water. **B:** Calculated velocities in vitreous. Solid lines correspond to a height of 20 µm and dashed lines to 10 µm. Purple curves show the thermal convection contribution, cyan curves show the thermophoretic contribution, and dark blue curves represent the total combined velocity. Compared to water, lower convection and thermophoresis velocities are observed in vitreous due to its higher viscosity.

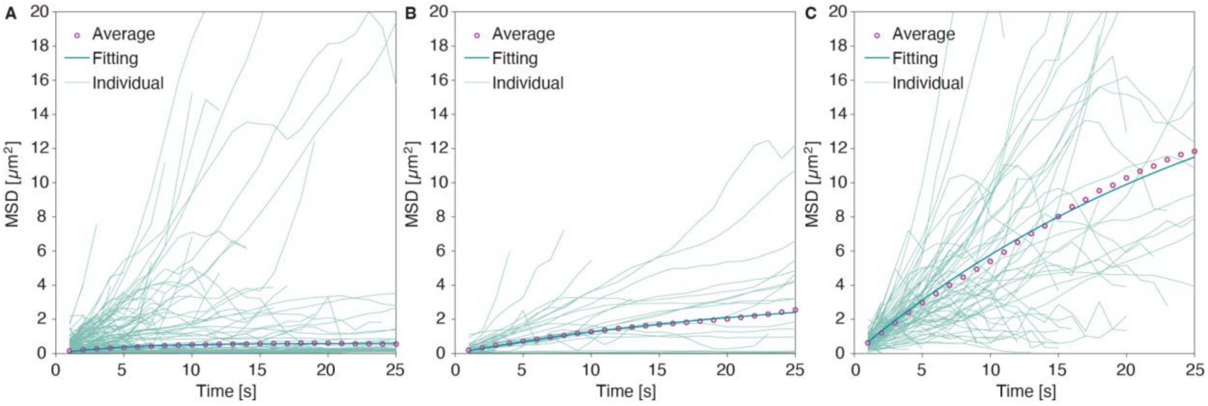


**Supplementary Figure S7. Motion of 520-nm PS nanoparticle (1/1000 (v/v)) in aged vitreous samples without ICG or laser irradiation.** Individual MSD traces obtained from MPT analysis are shown alongside exponential confinement model fits for samples analyzed at 0 (**A**), 4 (**B**), and 7 (**C**) days after extraction. The number of particle trajectories analyzed for each condition was 447, 53, and 102, respectively.

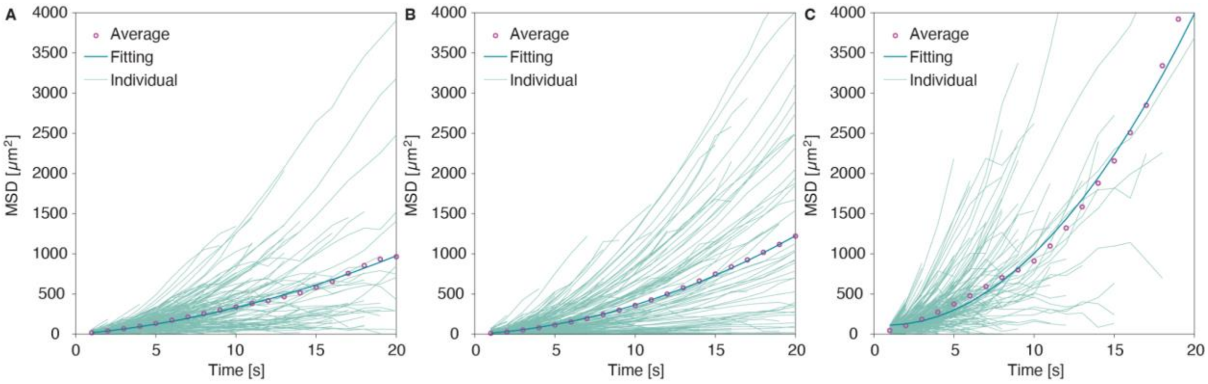


**Supplementary Figure S8. Photothermal transport of 520-nm PS nanoparticles (1/1000 (v/v)) in the presence of ICG (0.5 mg/mL) in aged vitreous samples upon laser irradiation (0.69 J/cm², 532 nm).** Individual mean squared displacement (MSD) traces from MPT analysis are shown for samples analyzed at 0 (**A**), 2 (**B**), and 8 (**C**) days after extraction, with corresponding quadratic model fits indicating active transport behavior. The number of particle trajectories analyzed for each condition was 327, 341, and 475, respectively.

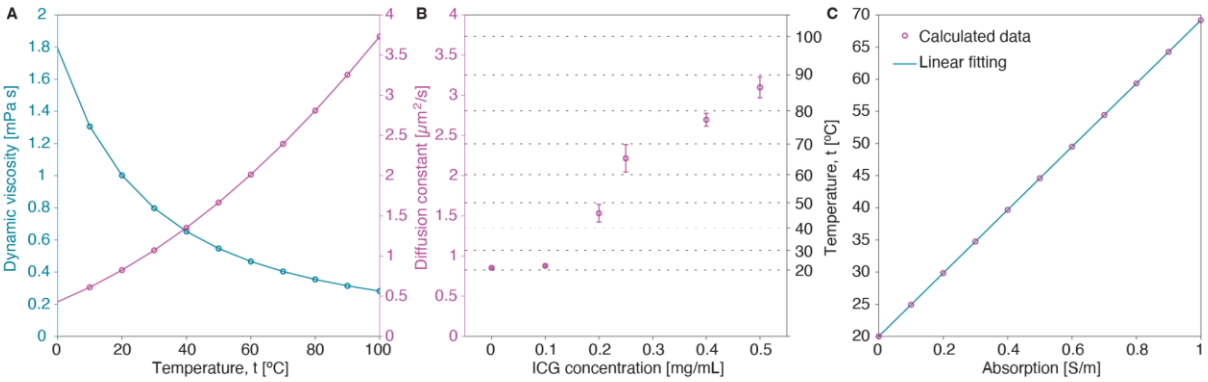


**Supplementary Figure S9. Photothermal heating and its impact on fluid viscosity and PS nanoparticle diffusion. A:** Temperature-dependent changes in dynamic viscosity of water (left Y-axis) and corresponding diffusion coefficients (right Y-axis) of 520-nm nanoparticles, highlighting the inverse relationship between temperature and viscosity and the resulting enhancement in diffusivity. **B:** Experimentally measured diffusion coefficients of 520-nm PS nanoparticles in water (1/5000 (v/v)) as a function of ICG concentration under laser irradiation (2.07 J/cm$^2$, 532 nm) shown in **Figure 1H** (see main text), resulting in elevated local temperatures (right Y-axis) and increased particle diffusivity (left Y-axis). **C:** Numerically-simulated temperature rise as a function of absorption coefficient, demonstrating a linear relationship between optical absorption and local heating under fixed laser power (100mW).

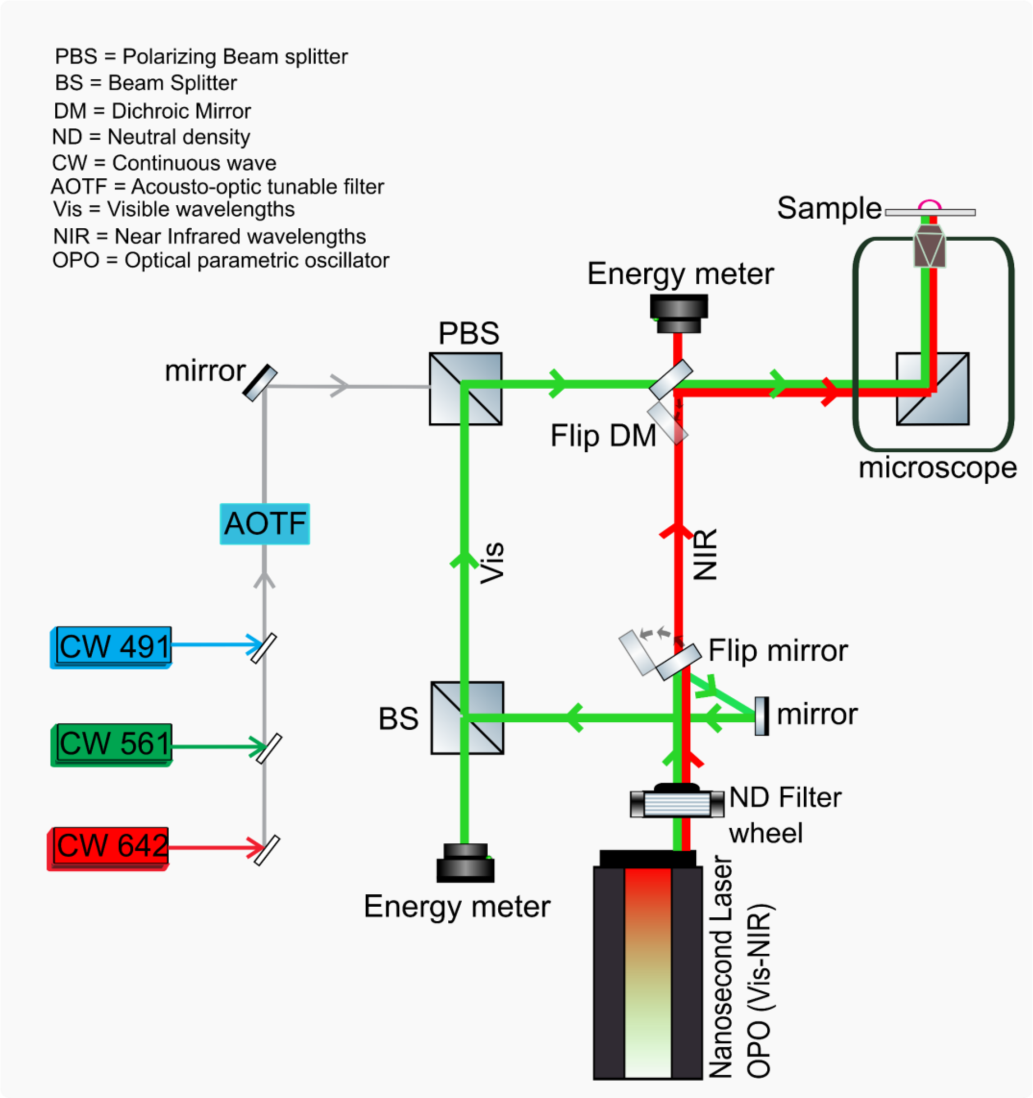


**Supplementary Figure S10. Schematic illustration of the nanosecond pulsed laser set-up (see Methods).**
