## Supplementary material for "Photothermal transport for guiding nanoparticles through the vitreous humor": Videos

### Slide 1
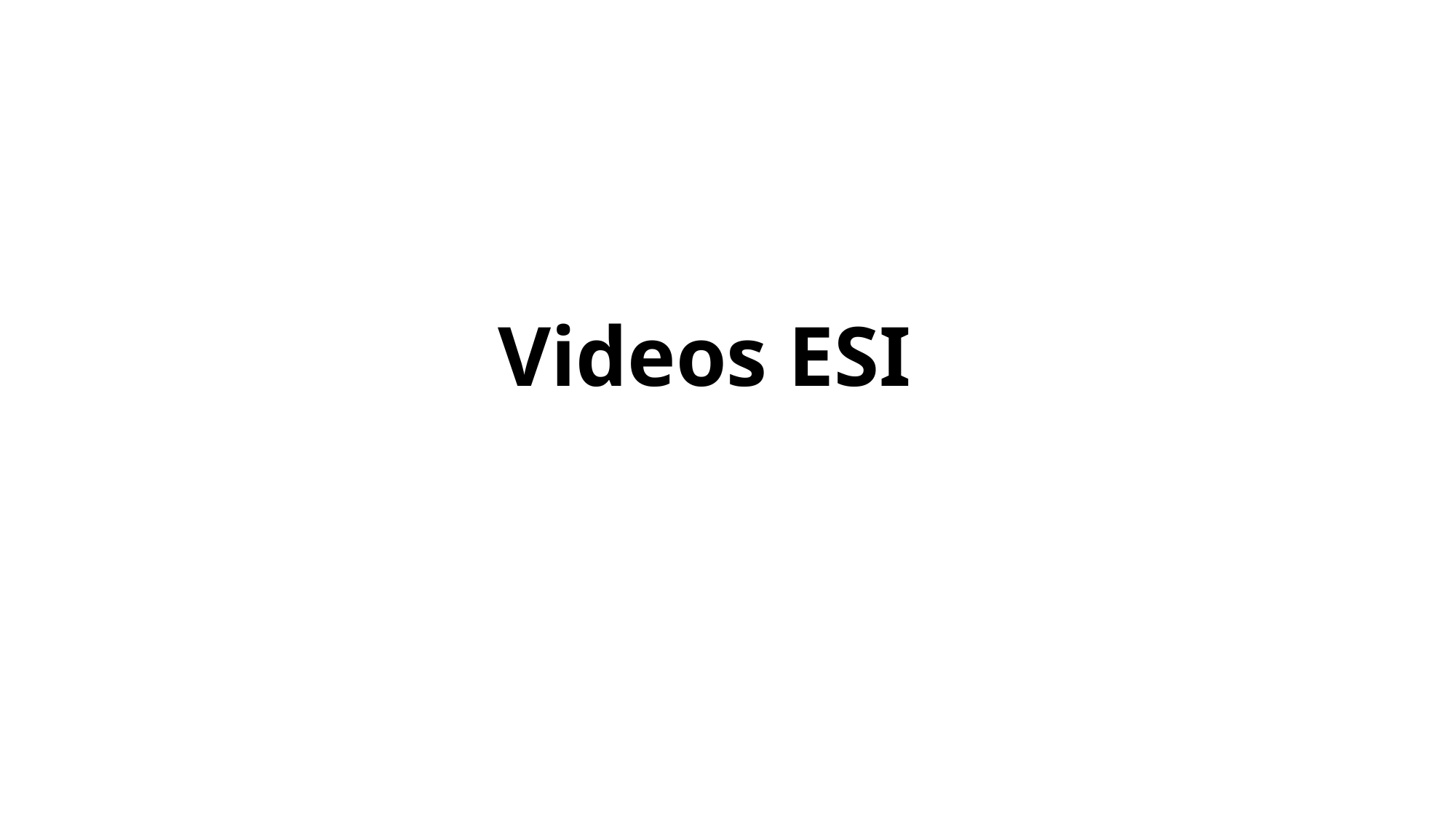

Videos ESI

### Slide 2
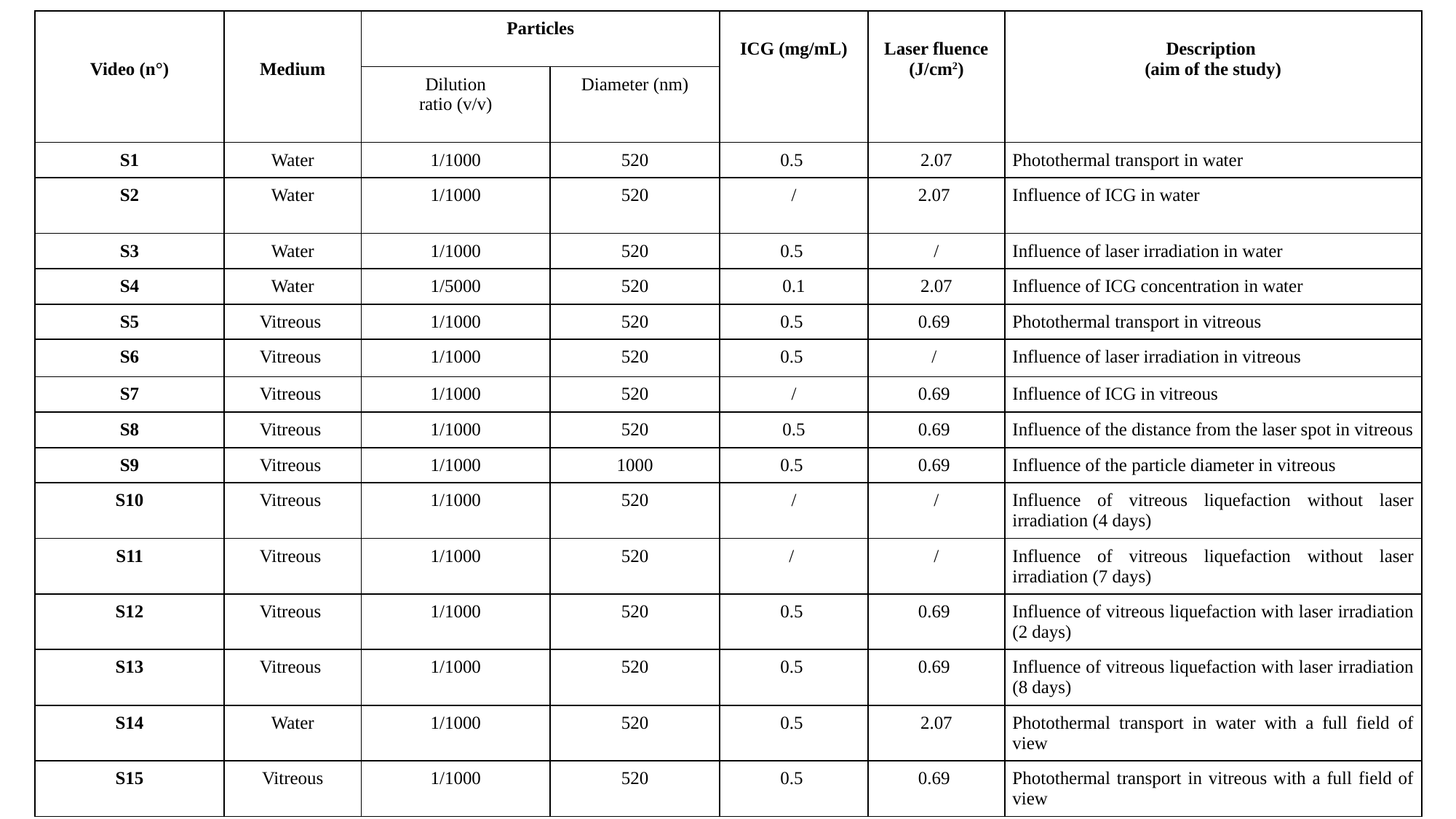

| Video (n°) | Medium | Particles | | ICG (mg/mL) | Laser fluence (J/cm^2^) | Description (aim of the study) |
| --- | --- | --- | --- | --- | --- | --- |
| | | Dilution ratio (v/v) | Diameter (nm) | | | |
| S1 | Water | 1/1000 | 520 | 0.5 | 2.07 | Photothermal transport in water |
| S2 | Water | 1/1000 | 520 | / | 2.07 | Influence of ICG in water |
| S3 | Water | 1/1000 | 520 | 0.5 | / | Influence of laser irradiation in water |
| S4 | Water | 1/5000 | 520 | 0.1 | 2.07 | Influence of ICG concentration in water |
| S5 | Vitreous | 1/1000 | 520 | 0.5 | 0.69 | Photothermal transport in vitreous |
| S6 | Vitreous | 1/1000 | 520 | 0.5 | / | Influence of laser irradiation in vitreous |
| S7 | Vitreous | 1/1000 | 520 | / | 0.69 | Influence of ICG in vitreous |
| S8 | Vitreous | 1/1000 | 520 | 0.5 | 0.69 | Influence of the distance from the laser spot in vitreous |
| S9 | Vitreous | 1/1000 | 1000 | 0.5 | 0.69 | Influence of the particle diameter in vitreous |
| S10 | Vitreous | 1/1000 | 520 | / | / | Influence of vitreous liquefaction without laser irradiation (4 days) |
| S11 | Vitreous | 1/1000 | 520 | / | / | Influence of vitreous liquefaction without laser irradiation (7 days) |
| S12 | Vitreous | 1/1000 | 520 | 0.5 | 0.69 | Influence of vitreous liquefaction with laser irradiation (2 days) |
| S13 | Vitreous | 1/1000 | 520 | 0.5 | 0.69 | Influence of vitreous liquefaction with laser irradiation (8 days) |
| S14 | Water | 1/1000 | 520 | 0.5 | 2.07 | Photothermal transport in water with a full field of view |
| S15 | Vitreous | 1/1000 | 520 | 0.5 | 0.69 | Photothermal transport in vitreous with a full field of view |

### Slide 3
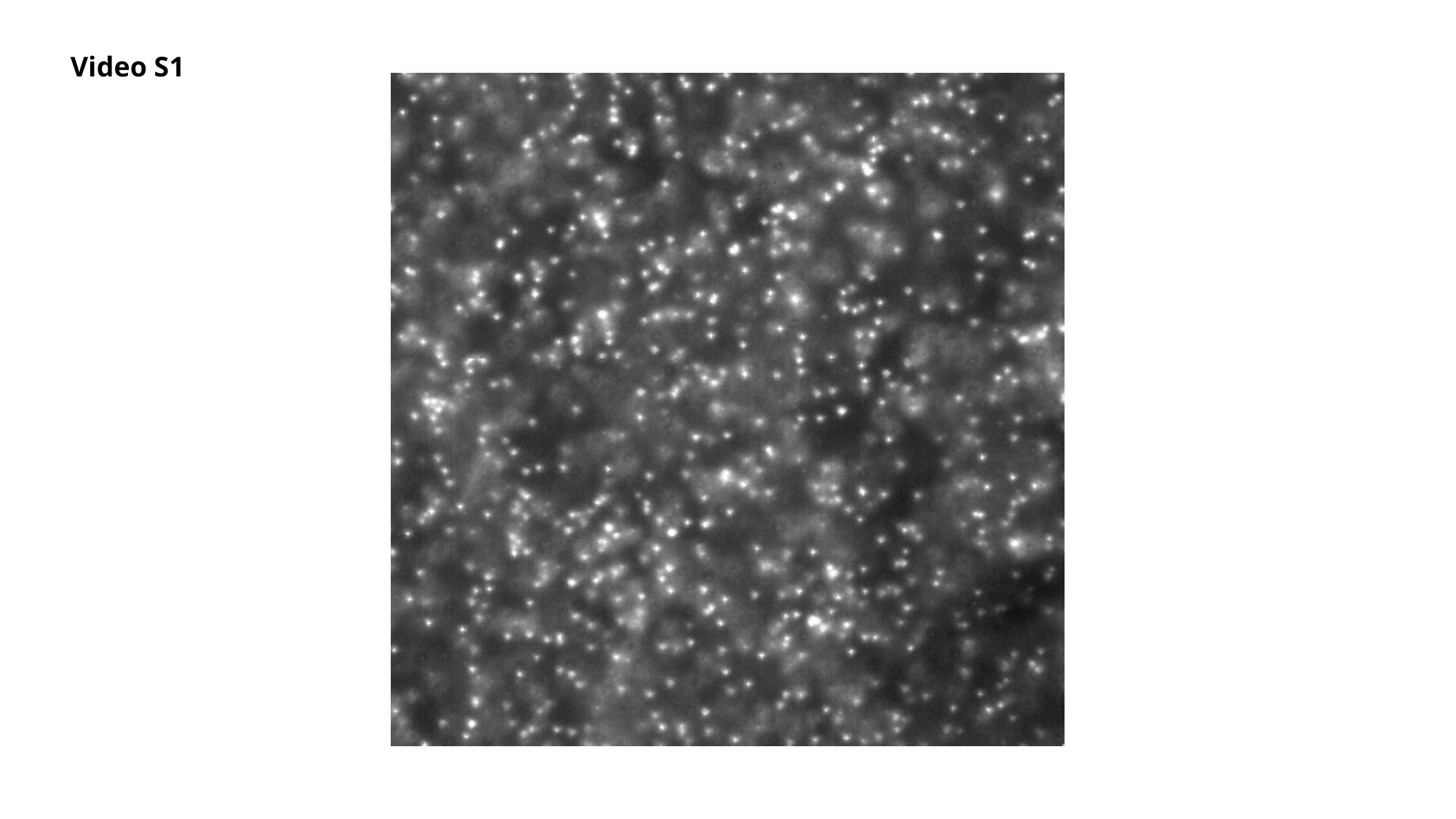

Video S1

### Slide 4
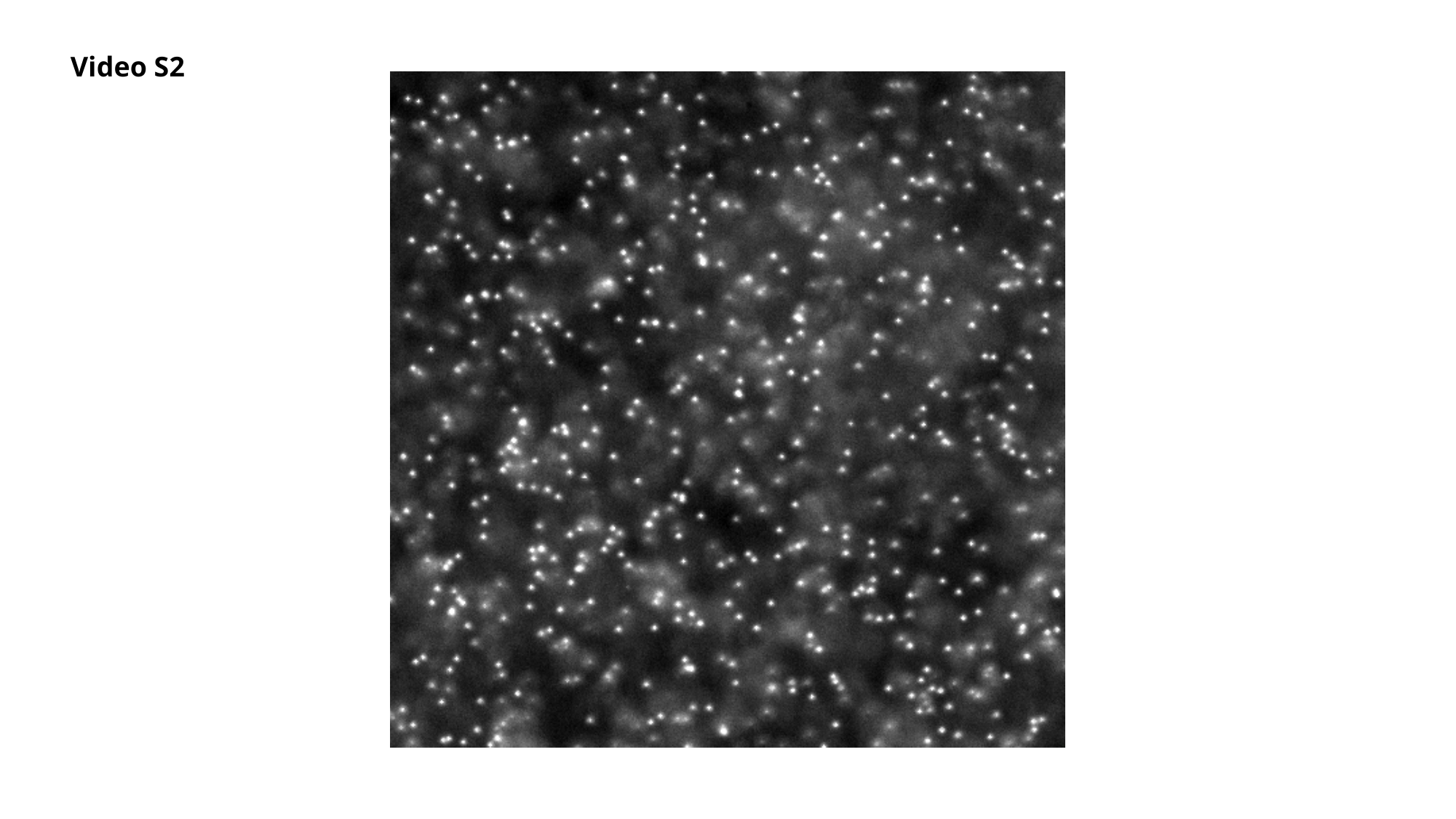

Video S2

### Slide 5
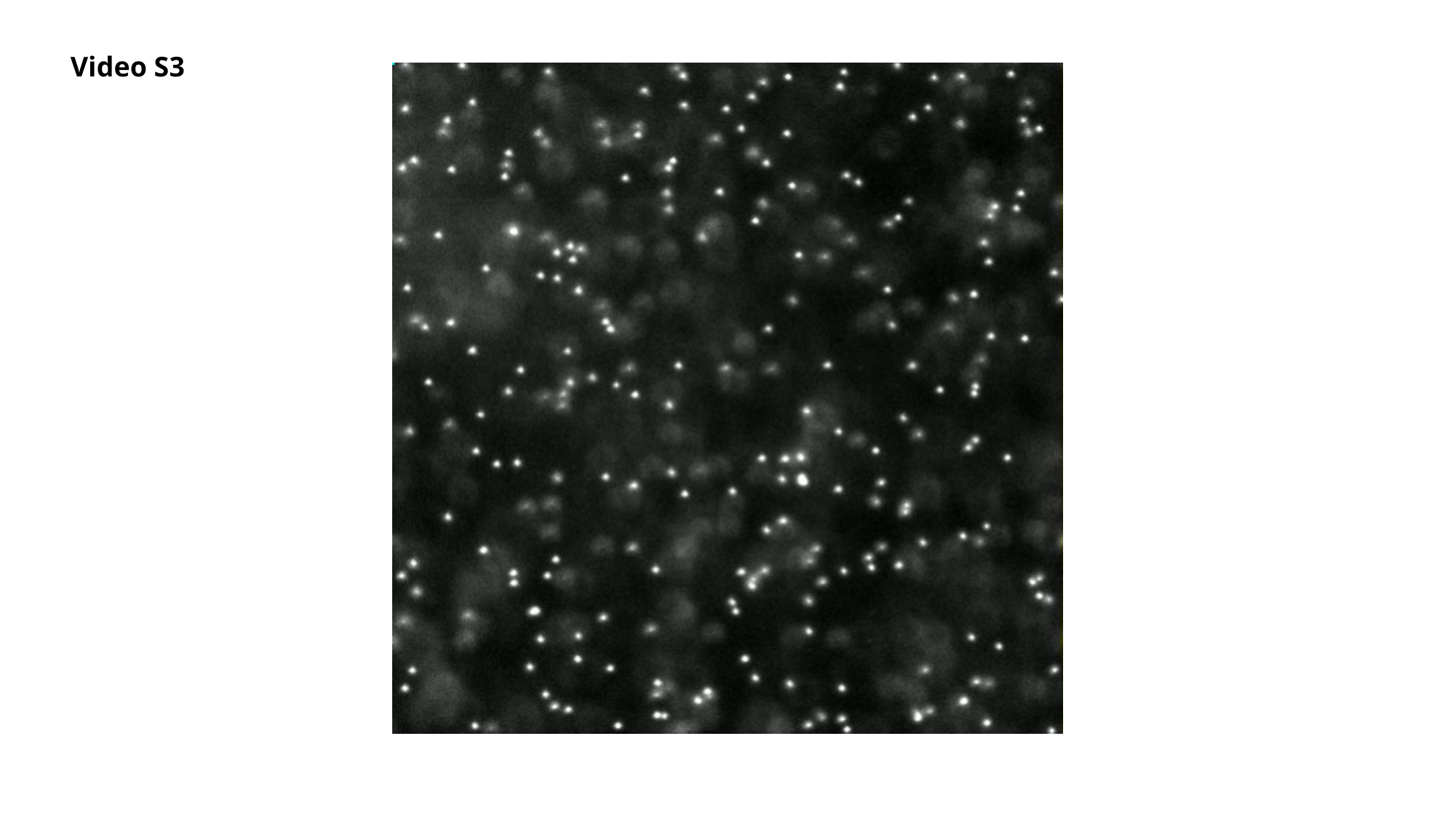

Video S3

### Slide 6
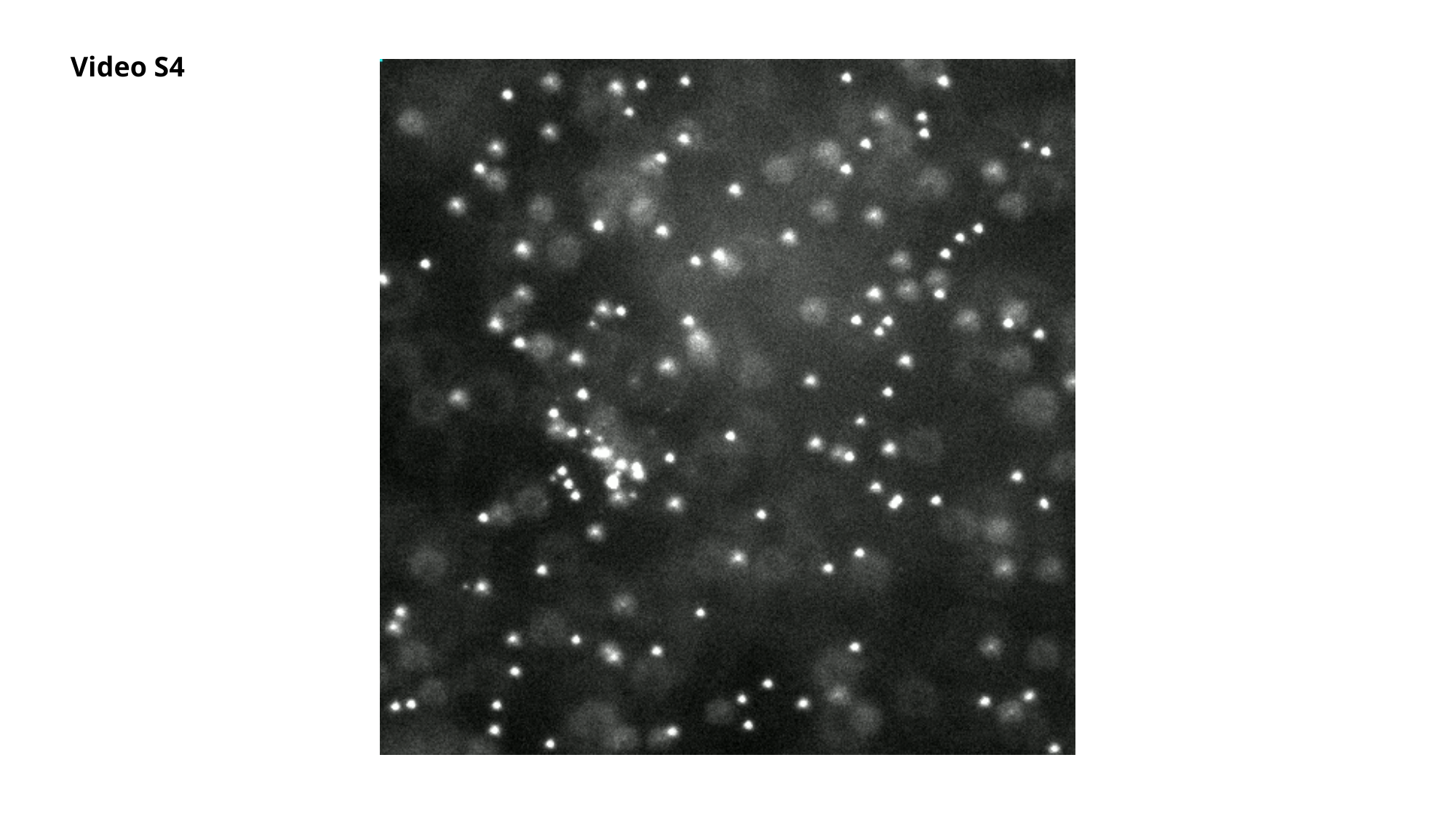

Video S4

### Slide 7
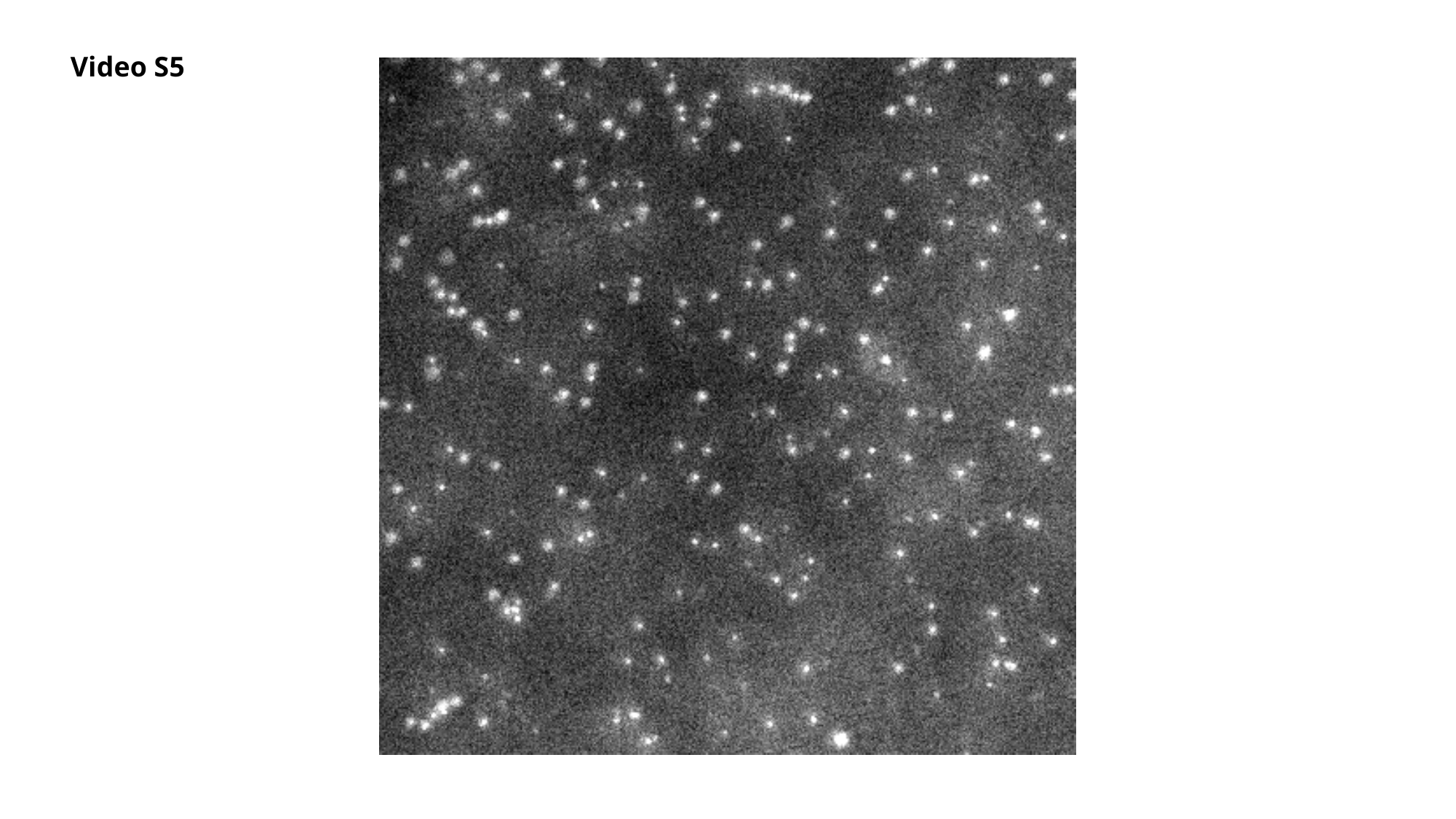

Video S5

### Slide 8
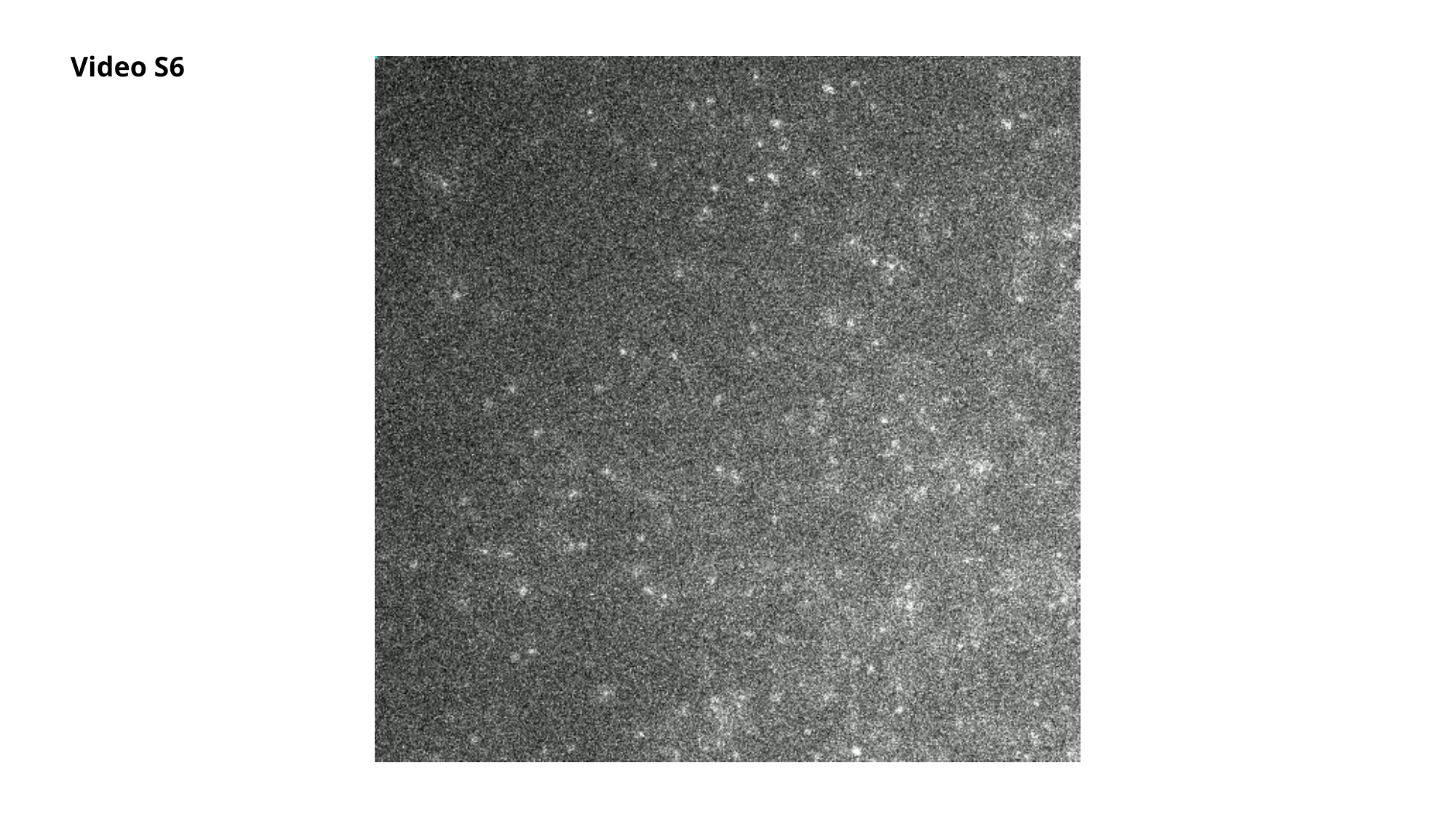

Video S6

### Slide 9
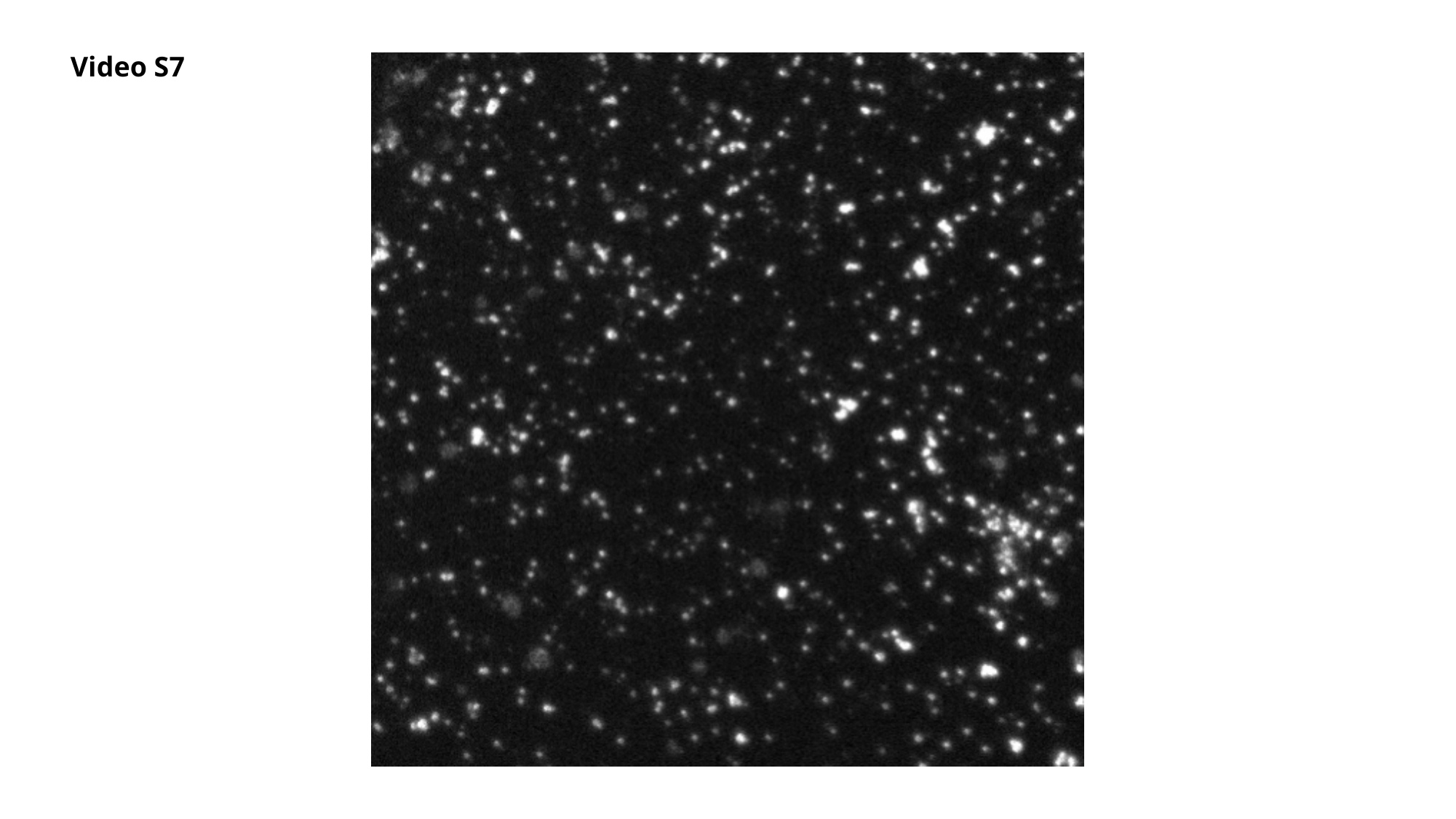

Video S7

### Slide 10
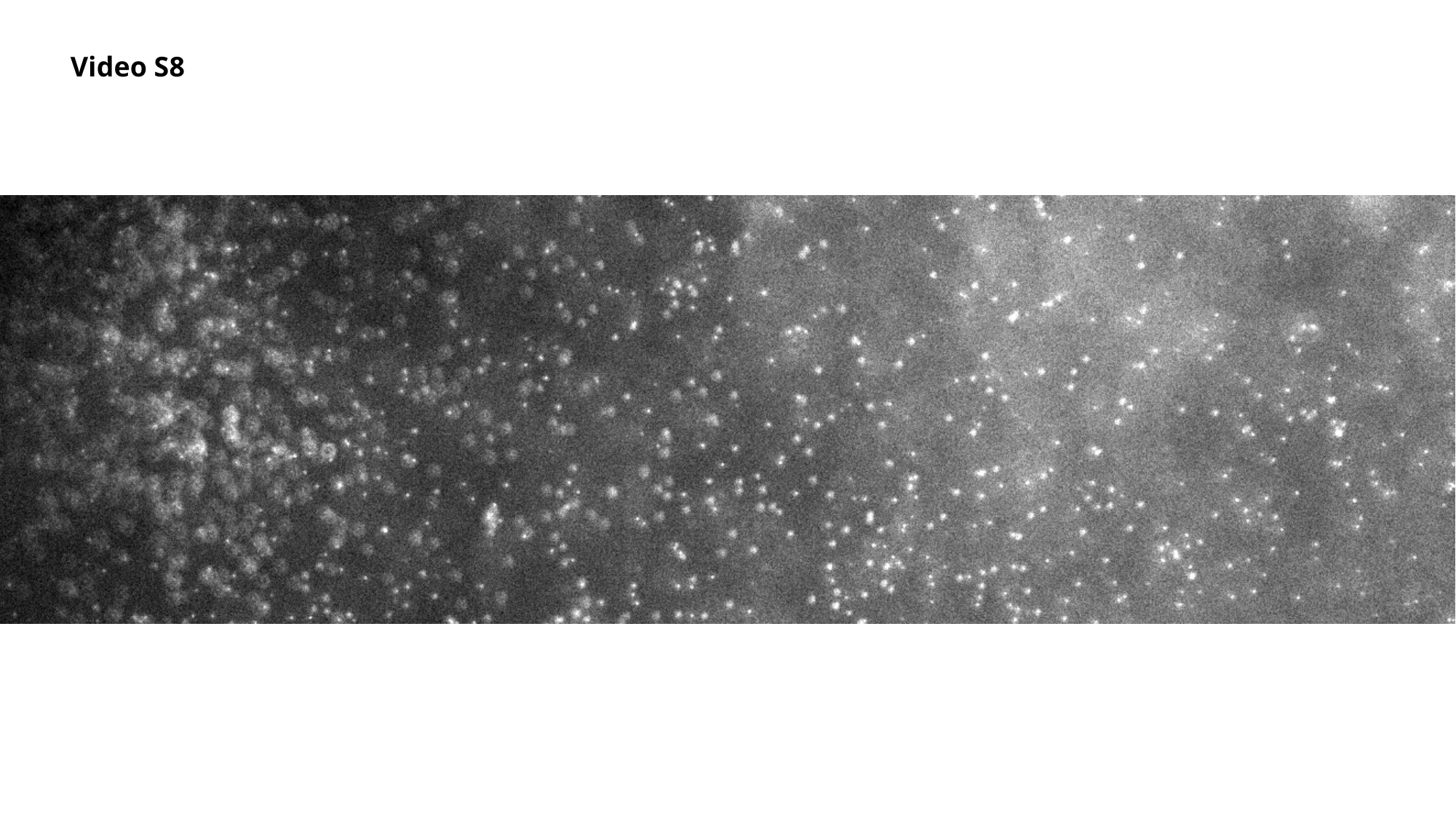

Video S8

### Slide 11
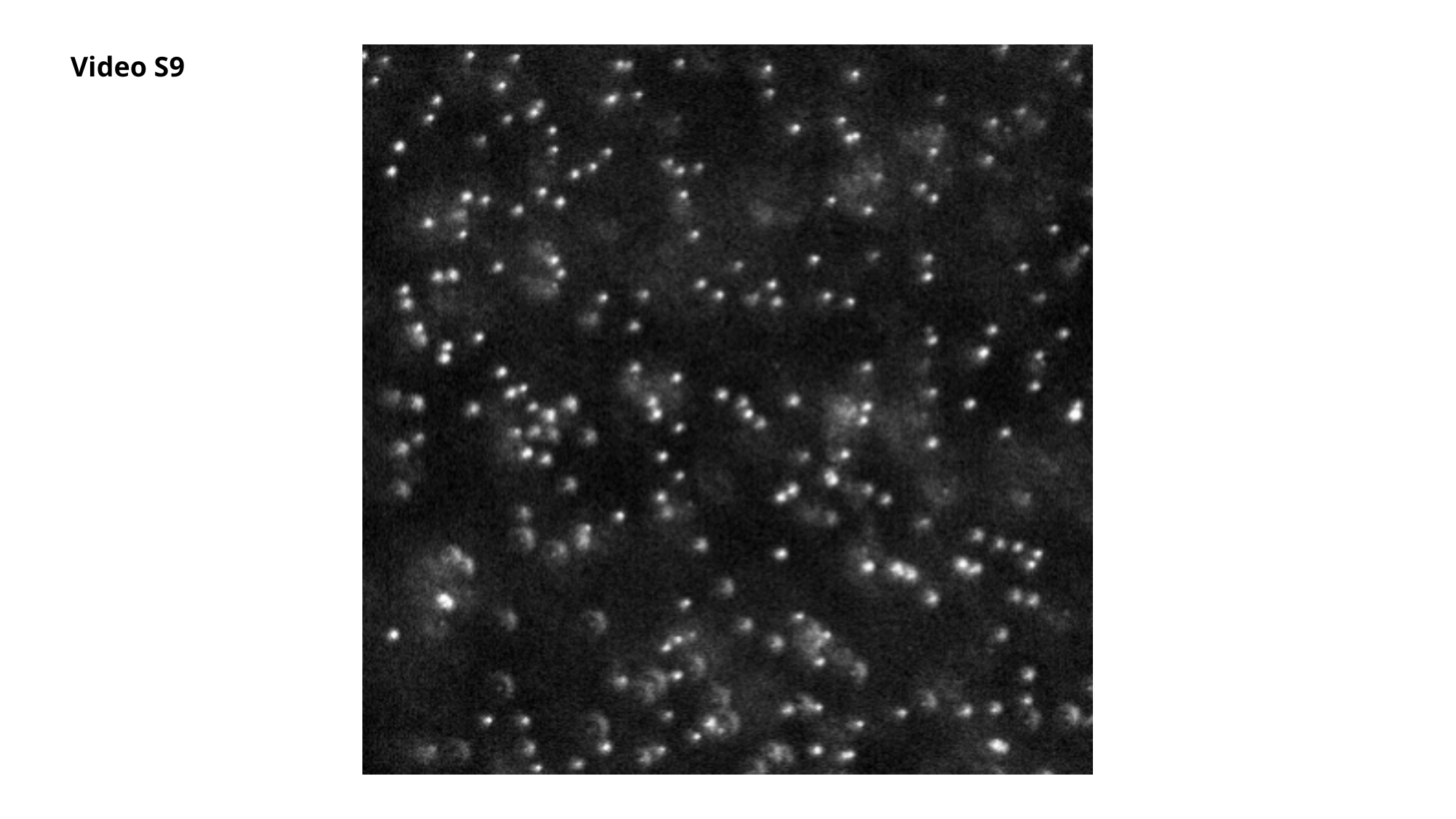

Video S9

### Slide 12
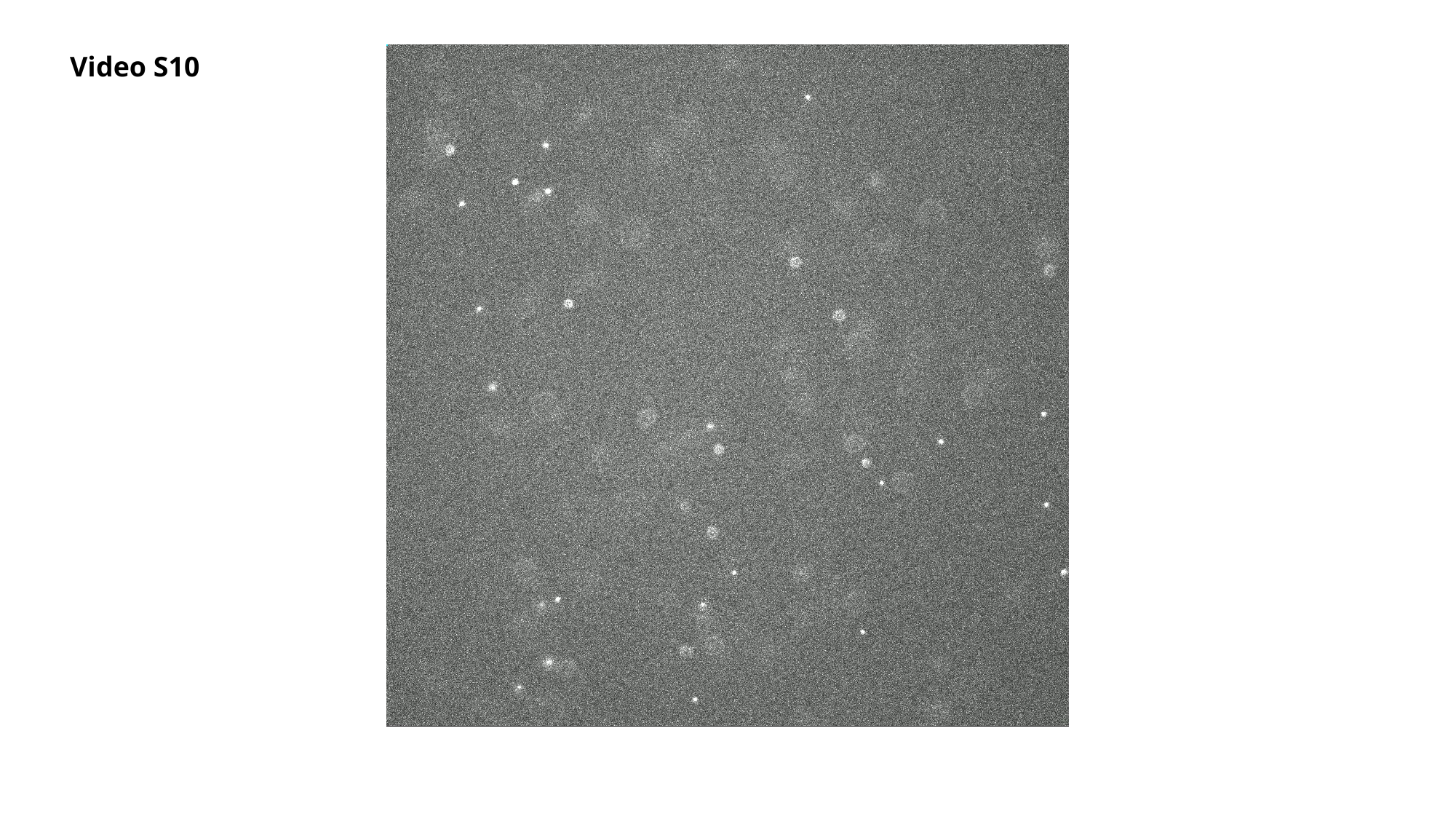

Video S10

### Slide 13
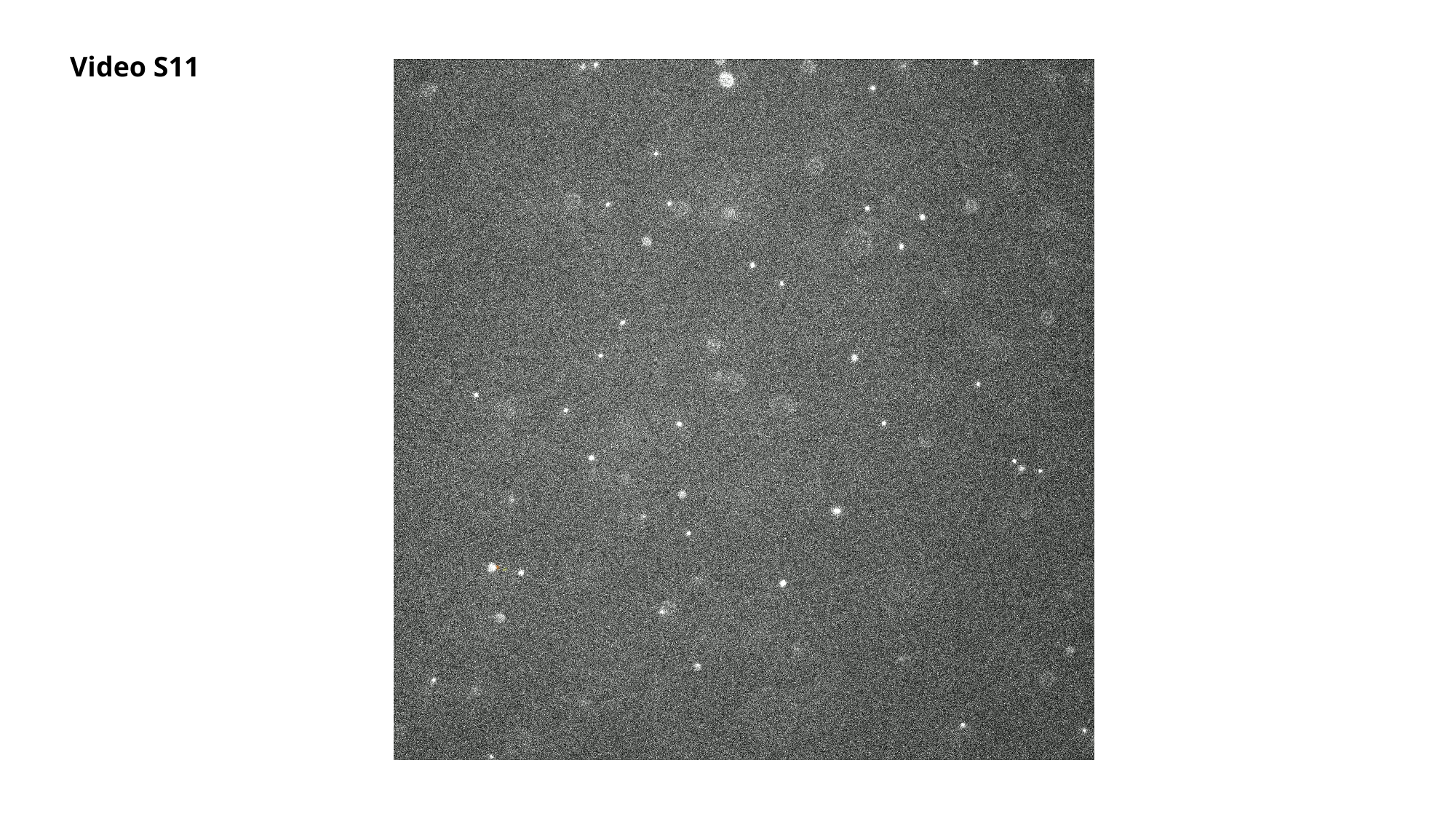

Video S11

### Slide 14
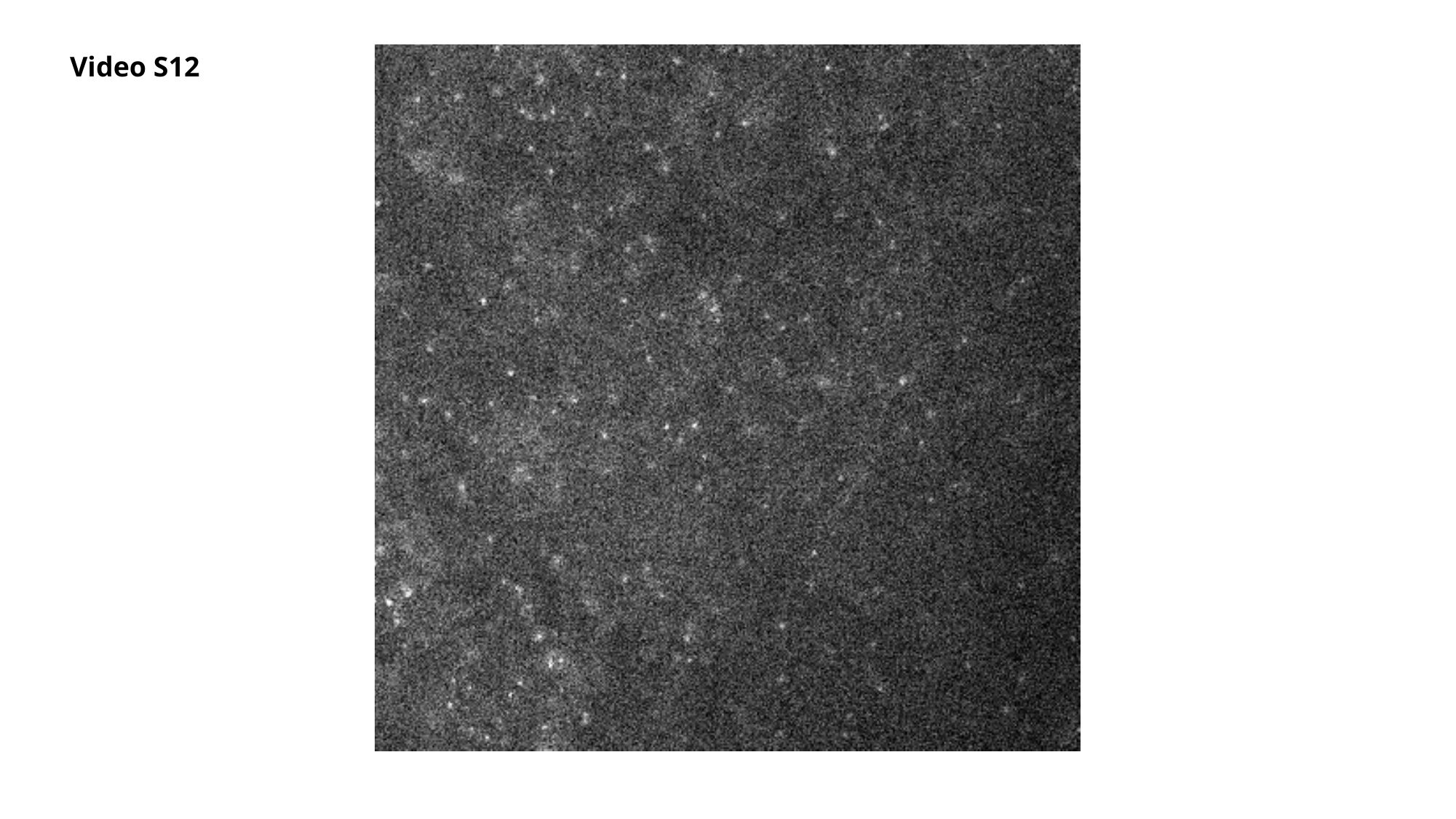

Video S12

### Slide 15
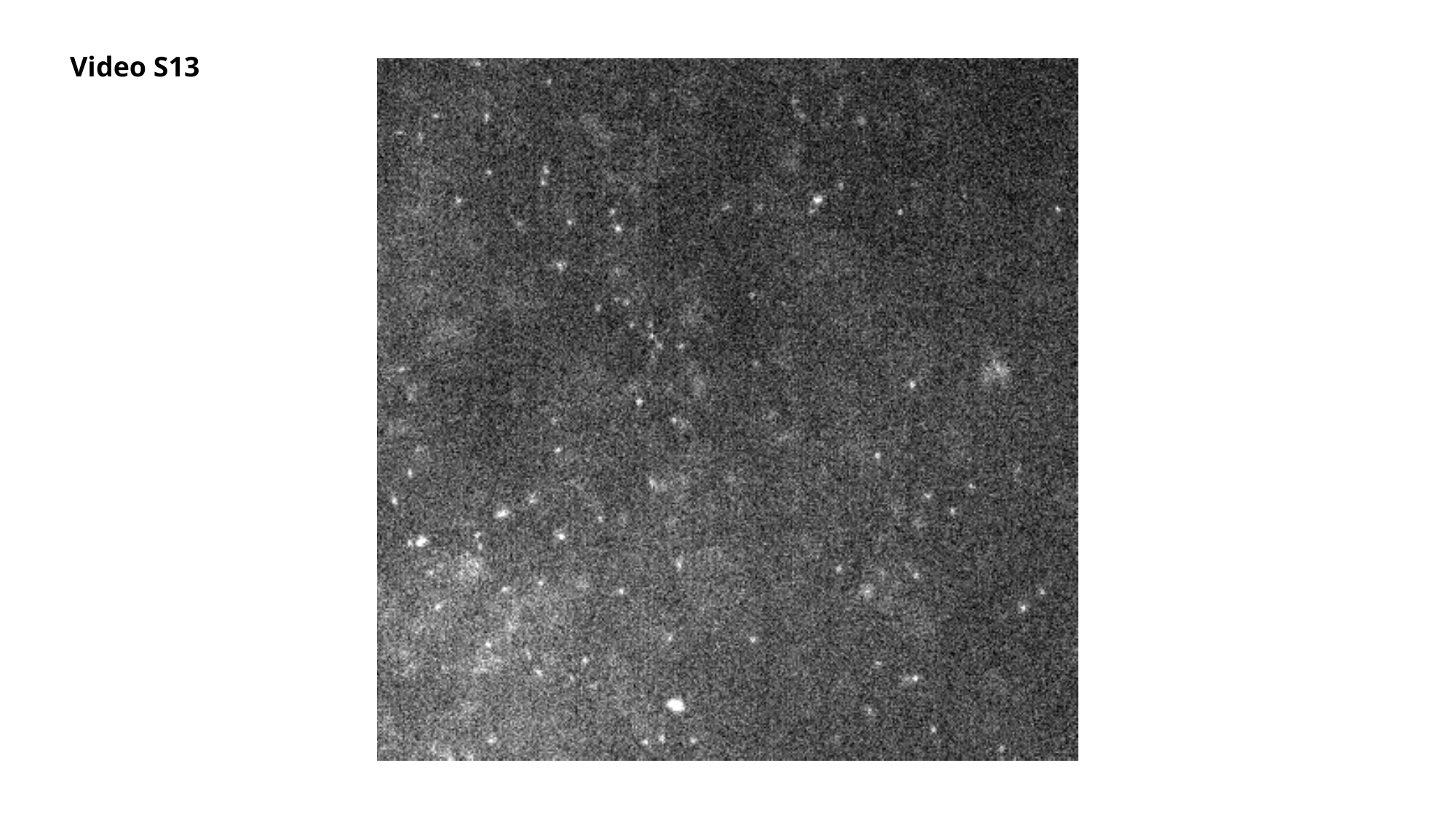

Video S13

### Slide 16
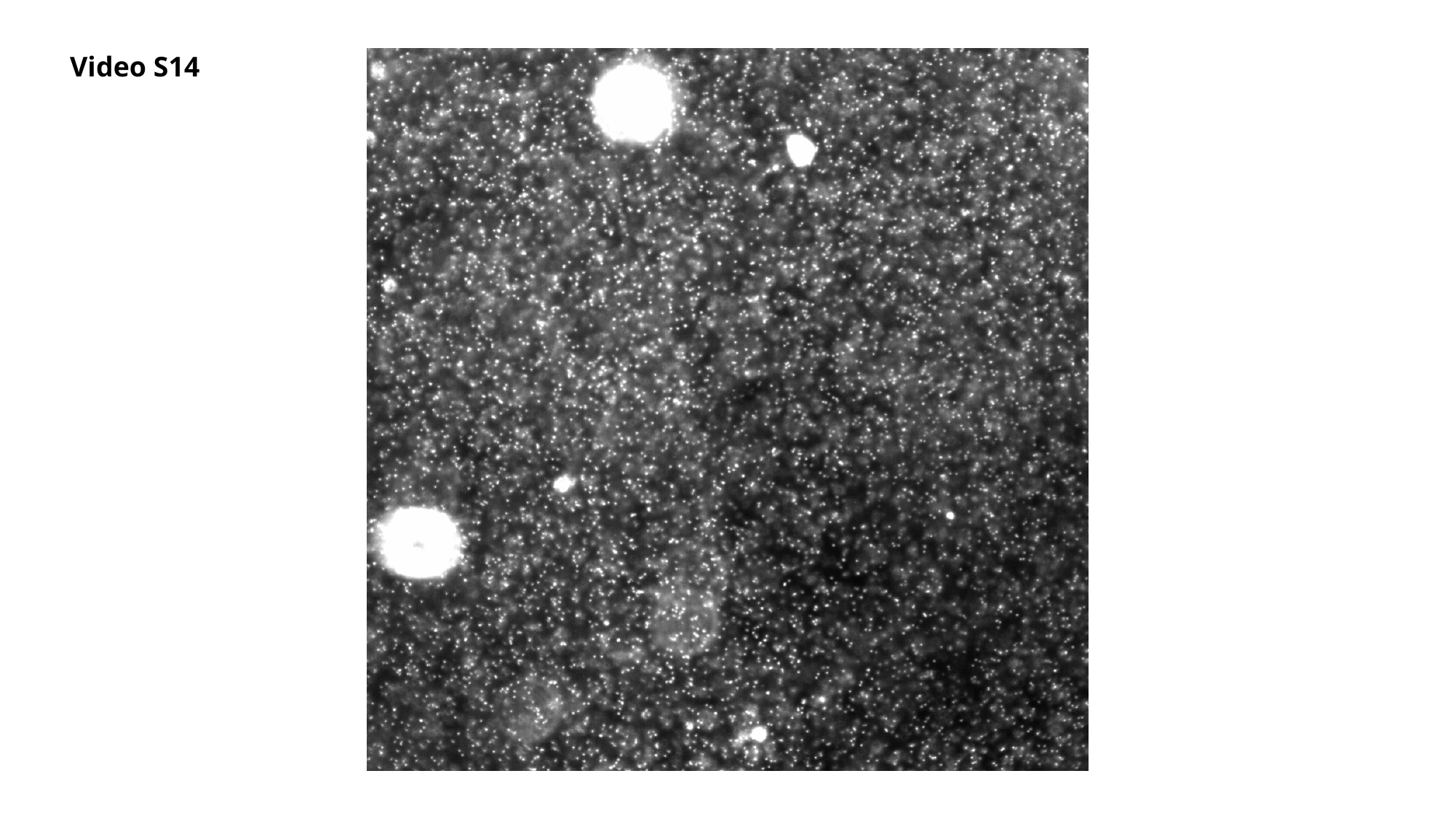

Video S14

### Slide 17
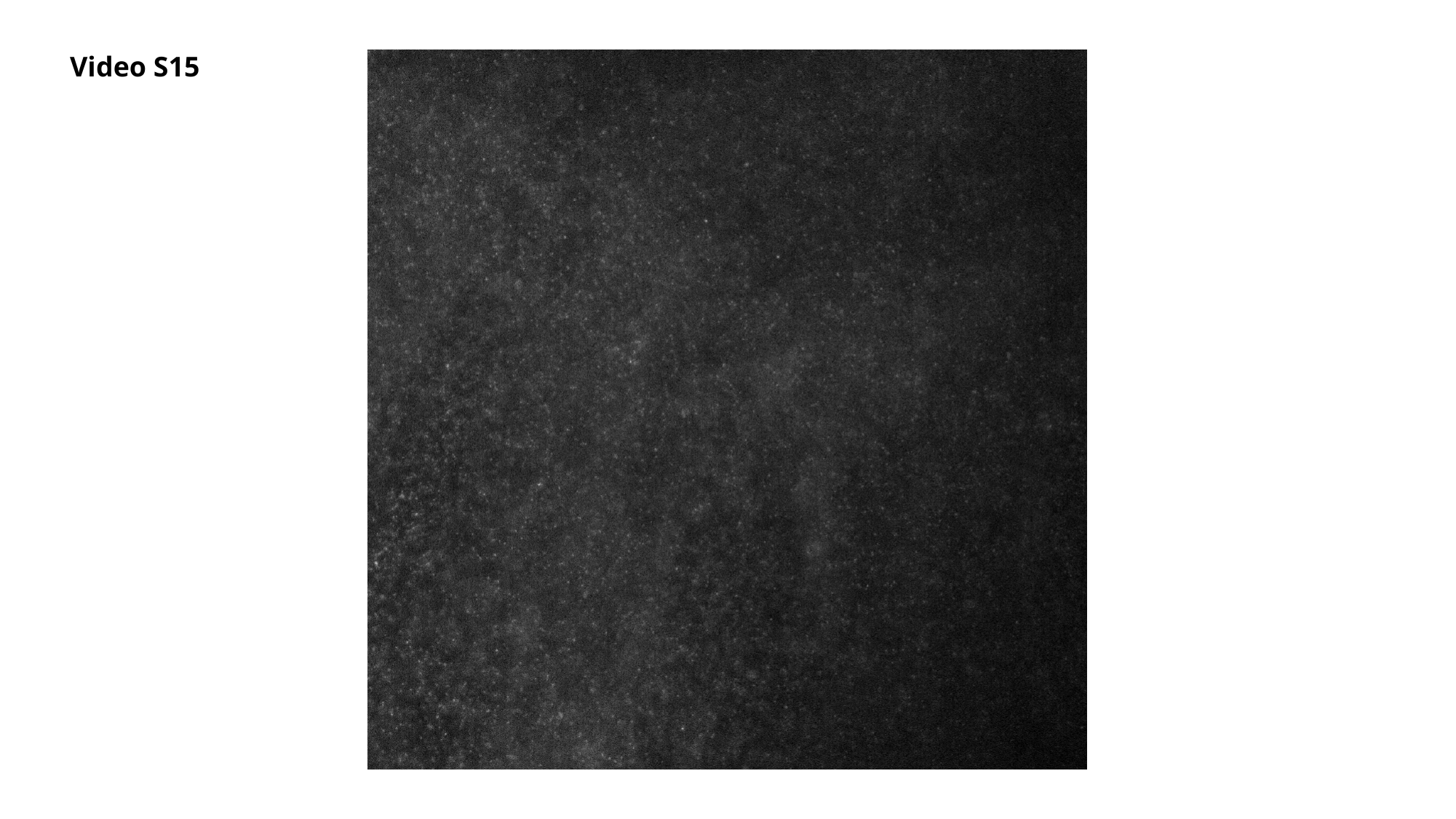

Video S15
